# Rrp1 translocase and ubiquitin ligase activities restrict the genome destabilising effects of Rad51 in fission yeast

**DOI:** 10.1101/2020.05.30.125286

**Authors:** Jakub Muraszko, Bilge Argunhan, Kentaro Ito, Gabriela Baranowska, Anna Barg-Wojas, Karol Kramarz, Yumiko Kurokawa, Hiroshi Iwasaki, Dorota Dziadkowiec

**Affiliations:** Faculty of Biotechnology, University of Wrocław, Poland; Cell Biology Center, Institute of Innovative Research, Tokyo Institute of Technology, Japan; Institut Curie, Centre National de la Recherche Scientifique, Orsay, France; Center for Frontier Research, National Institute of Genetics, Japan

## Abstract

Rad51 is the key protein in homologous recombination DNA repair and has important roles during DNA replication. Auxiliary factors regulate Rad51 activity to facilitate productive, and prevent inappropriate, recombination that could lead to genome instability. Previous genetic analyses identified a function for Rrp1 (a member of the Rad5/16-like group of SWI2/SNF2 translocases) in counteracting Rad51 activity, shared with the Rad51 mediator Swi5-Sfr1 and the Srs2 anti-recombinase. Here, we show that Rrp1 overproduction alleviates the toxicity associated with excessive Rad51 activity in a manner dependent on Rrp1 ATPase domain. Purified Rrp1 binds to DNA and has a DNA-dependent ATPase activity. Importantly, Rrp1 directly interacts with Rad51 and removes it from double-stranded DNA, confirming that Rrp1 is a translocase capable of modulating Rad51 activity. Additionally, we demonstrate that Rrp1 possesses E3 ubiquitin ligase activity with Rad51 as a substrate, suggesting that Rrp1 regulates Rad51 in a multi-tiered fashion.

## Introduction

Homologous recombination (HR) is a highly conserved pathway that is involved in the repair of DNA double-strand breaks (DSBs), and many key HR proteins have a critical role during DNA replication (Kolinjivadi et al., 2017). During DSB repair, the Rad51 recombinase forms a nucleoprotein filament on single-stranded DNA (ssDNA) that catalyses strand invasion into intact homologous double-stranded DNA (dsDNA) (Kowalczykowski, 2015; Symington, 2002). Rad51 is aided by a group of proteins called recombination mediators. The main mediator in yeasts, Rad52, facilitates Rad51 loading onto replication protein A (RPA)-coated ssDNA (Krogh and Symington, 2004; Raji and Hartsuiker, 2006). In human cells, the tumour suppressor protein BRCA2 fulfils an equivalent function during HR, recruiting Rad51 onto RPA-coated ssDNA and stabilising presynaptic filaments (Liu and Heyer, 2011). Additionally, human cells contain five Rad51 paralogs (RAD51B, RAD51C, RAD51D, XRCC2 and XRCC3) that influence HR, and these factors are thought to stimulate Rad51 activity, reviewed in (Prakash et al., 2015). In the fission yeast *Schizosaccharomyces pombe*, two auxiliary factor complexes have been shown to promote Rad51-dependent DNA repair: Rad55-Rad57 and Swi5-Sfr1 (Akamatsu et al., 2007; Argunhan et al., 2020), both of which are conserved in humans (Argunhan et al., 2017; Yuan and Chen, 2011). We previously identified another complex, Rrp1-Rrp2, that acts in a Swi5-Sfr1-dependent sub-pathway of HR in the replication stress response (Dziadkowiec et al., 2009, 2013).

Replication forks frequently stall at specific sites in the genome, such as repetitive DNA sequences, DNA lesions resulting from exogenous damage or sites of DNA-protein association (Carr et al., 2011), and Rad51 has multiple important functions at such arrested forks. First, Rad51 binding stabilizes replication forks by protecting them from nuclease degradation (Hashimoto et al., 2010; Schlacher et al., 2011). Second, Rad51 participates in replication fork reversal, a global mechanism to stabilise forks and protect them from breakage, and stimulates the fork regression activity of Rad54, the SWI2/SNF2-like translocase, by inhibiting fork restoration (Bugreev et al., 2011; Zellweger et al., 2015). Several SNF2-family DNA translocases, such as SMARCAL1, ZRANB3 and HLTF, are able to remodel replication forks (Achar et al., 2015; Bétous et al., 2012; Kile et al., 2015; Vujanovic et al., 2017), and Rad51 is proposed to cooperate in this process by driving the equilibrium of the reaction toward fork reversal (Bhat and Cortez, 2018). Interestingly, fork regression is not dependent on the presence of the BRCA2 mediator (Mijic et al., 2017). Finally, when the replication fork is inactivated or converted to a DSB by MUS81-dependent nucleolytic cleavage, the strand exchange activity of Rad51 promotes HR-dependent reconstitution of replication (Lambert et al., 2005; McGlynn and Lloyd, 2002; Pepe and West, 2014).

Rad51 filament formation must be tightly regulated because inappropriate, excessive, or untimely recombination (especially at replication forks or repeat sequences) can lead to deleterious effects including loss of heterozygosity and chromosome rearrangements that are hallmarks of cancer in higher organisms (Sung and Klein, 2006). Many helicases, such as Srs2 and Sgs1 in yeast, and their respective mammalian orthologues PARI and BLM, as well as FANCJ and RECQ5, have been implicated in regulating the stability of Rad51 filaments formed on ssDNA. This ensures that the HR-mediated DSB repair process is reversible and can proceed along multiple pathways, making it both flexible and robust, reviewed in (Heyer, 2015). Recently, RADX has been found to antagonise Rad51 binding to ssDNA specifically at replication forks, where it regulates the balance between Rad51 fork protection, fork reversal and its role in DSB repair (Bhat et al., 2018; Dungrawala et al., 2017).

Rad51 must not only be able to form filaments on ssDNA but also bind to dsDNA tracts in order to carry out its multiple functions in HR repair and at replication forks, and this process is also stringently regulated. The SWI2/SNF2-like translocase Rad54 dissociates Rad51 from dsDNA in both yeast and human cells (Mason et al., 2015; Solinger et al., 2002; Wright and Heyer, 2014), allowing the repair synthesis by DNA polymerases, necessary for the completion of HR. Rad54 is activated in G2, and does not remove Rad51 from stalled replication forks (Spies et al., 2016). Another complex, MMS22L-TONSL, has been shown in human cells to limit Rad51 binding to dsDNA and stimulate HR-mediated restart of arrested replication forks (Piwko et al., 2016). Importantly, other SWI2/SNF2-like translocases, Rdh54 and Uls1, cooperate with Rad54 in *Saccharomyces cerevisiae* not only to antagonise Rad51 binding to dsDNA during HR, but also to counteract its toxic accumulation on undamaged chromatin (Chi et al., 2006, 2011; Shah et al., 2010). RAD54L and RAD54B in humans also prevent the genome-destabilising consequences of excessive Rad51 binding to dsDNA (Mason et al., 2015). It should also be noted that the binding of Rad51 to dsDNA renders the dsDNA inaccessible to the Rad51-ssDNA filament and thus acts as a barrier to HR itself (Sung and Robberson, 1995).

Interestingly, the Rad51 paralog RAD51C has been shown to prevent proteasomal degradation of Rad51 in human cells, especially after DNA damage (Bennett and Knight, 2005), suggesting that Rad51 can also be regulated by ubiquitylation. Recently, Rad51 was found to be poly-ubiquitylated by the E3 ubiquitin ligase RFWD3 in a process stimulated by DNA damage (Inano et al., 2017). Rad51 ubiquitylation decreases Rad51 binding to ssDNA and leads to its proteasomal degradation, while also stimulating chromatin loading of Rad54. It has therefore been proposed that Rad51 ubiquitylation promotes its removal from sites of DNA damage and is necessary for completion of HR DNA repair (Inano et al., 2017).

Regulation of Rad51 by Fbh1, an F-box helicase and E3 ubiquitin ligase, is more complex and involves both activities of this protein. In *S. pombe*, Fbh1 acts as a translocase and disrupts Rad51 ssDNA filaments, thereby regulating the outcome of the HR reaction. Additionally, the SCF^Fbh1^ E3 complex can ubiquitylate Rad51 *in vitro* and this modification was found to be necessary for depletion of Rad51 in stationary-phase cells (Tsutsui et al., 2014). Human Rad51 is also monoubiquitylated *in vitro* by SCF^FBH1^ but this does not result in the protein’s turnover by proteolysis (Chu et al., 2015). Instead, the authors propose that during replication stress, FBH1 translocase displaces Rad51 from ssDNA and modifies it to prevent Rad51 from reloading, thus restricting untimely HR at the replication fork.

Many proteins with helicase, translocase and/or ubiquitin ligase activities have been found to regulate Rad51 activity, underscoring the importance of this process. Here, we show that Rrp1, an orthologue of *S. cerevisiae* Uls1, belonging to the unique RING-domain-containing Rad5/16-like group of SWI2/SNF2 translocases, ubiquitylates Rad51 and is able to displace it from dsDNA. We propose that these translocase and ubiquitin ligase activities allow Rrp1 to counteract the genotoxic effects of excessive Rad51 binding to undamaged chromatin.

## Results

### Rrp1 counteracts *rad51*^+^ overexpression-induced toxicity

Previous studies have shown that *rad51*^+^ overexpression in *S. pombe* results in a severe growth defect (Kim et al., 2001). It has been demonstrated that Rad51 overproduction leads to its excessive accumulation on chromatin and has a negative effect on cell growth and chromosome stability that is aggravated in mutants devoid of SWI2/SNF2-related translocases: RAD54L and RAD54B in humans (Mason et al., 2015) and Rdh54, Rad54, and Uls1 in *S. cerevisiae* (Shah et al., 2010). Two *ULS1* orthologues have been identified in *S. pombe* (Dziadkowiec et al., 2009), so we set out to examine their functional interaction with Rad51. Using an *nmtP3-GFP-rad51* strain, where the *rad51*^+^ gene is expressed from a strong promoter that is induced in media lacking thiamine, we confirmed that induction of *rad51*^+^ expression resulted in a severe growth defect (Figure 1A,B) and loss of viability (Figure 1C). Importantly, this was further exacerbated by deletion of *rrp1*^+^, but not *rrp2*^+^. Growth inhibition of the *nmtP3-GFP-rad51 rrp1*Δ strain was visible even on YES medium (Figure 1A,B), where gene expression from the *nmtP3* promoter is very limited, indicating that even a mild increase in Rad51 protein levels is toxic when the Rrp1 translocase is absent.

**Figure 1.**
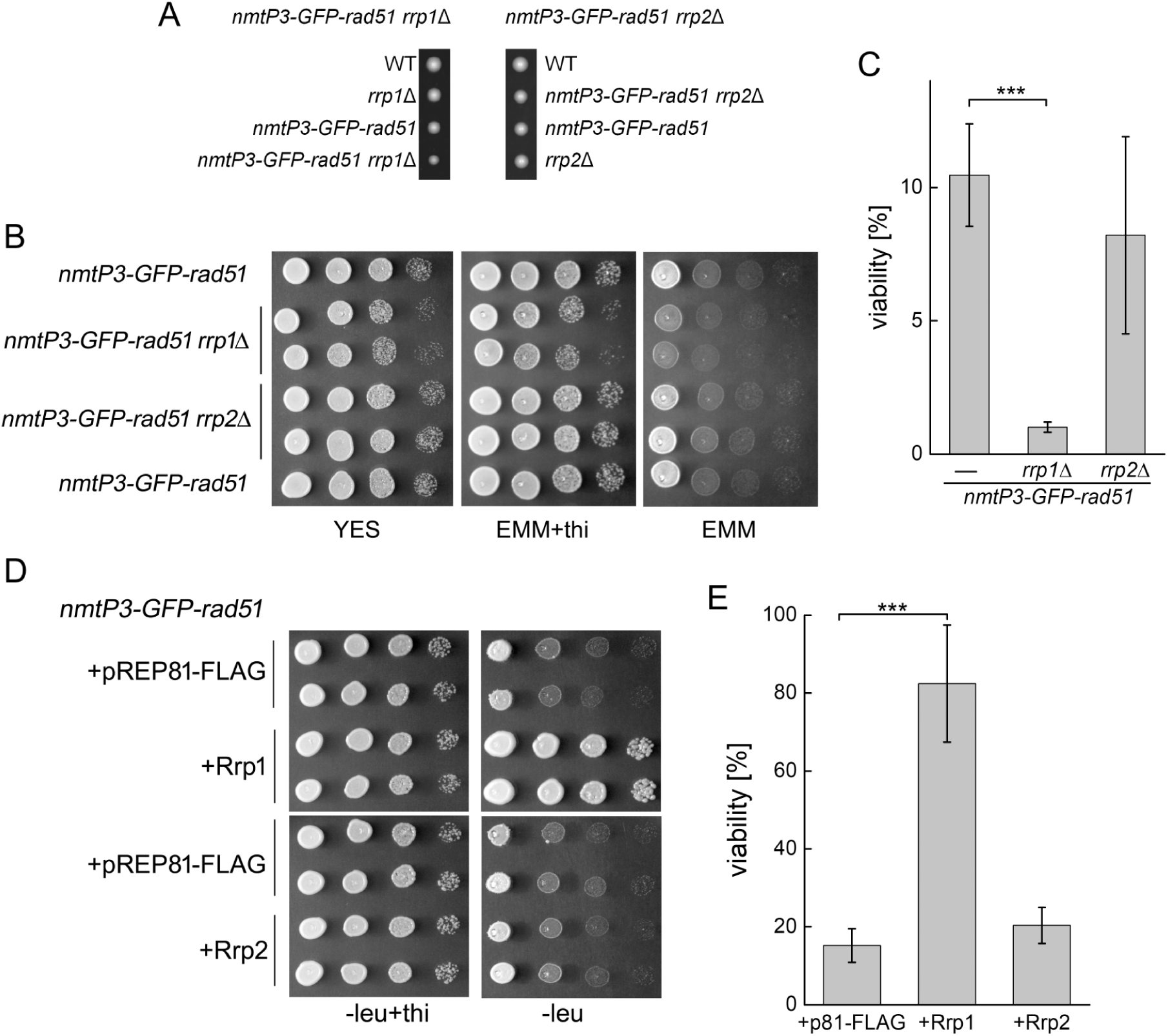
Rrp1 is important for viability in cells over-expressing *rad51*^+^. (A) The *nmtP3-GFP-rad51 rrp1*Δ strain is viable but exhibits a severe slow-growth phenotype, as determined by tetrad analysis. (B) Deletion of *rrp1*^+^, but not *rrp2*^+^, exacerbates the growth defect induced by *rad51*^+^ overexpression. Growth of the indicated strains under conditions where *rad51*^+^ overexpression was induced (EMM) or repressed (EMM +thiamine) was examined by spot test analysis. (C) Viability loss caused by *rad51*^+^ overexpression is greater in *rrp1*Δ cells. Strains were grown in the presence and absence of thiamine and the percentage of surviving cells was determined. The experiment was repeated twice. (D) Spot test analysis demonstrating that overexpression of *rrp1*^+^, but not *rrp2*^+^, suppresses the growth defect caused by Rad51 overproduction. (E) Simultaneous *rrp1*^+^ overexpression reverses viability loss in cells overexpressing *rad51*^+^, determined by comparing the growth of transformants in the presence and the absence of thiamine. The experiment was repeated with at least three independent transformants. Error bars represent the standard deviation about the mean values. Student’s t-test was used to calculate the *p*-value (*** *p*-value ≤ 0.001).

Due to the severe growth defect of the *nmtP3-GFP-rad51 rrp1*Δ strain, as well as the rapid generation and subsequent proliferation of faster-growing suppressors, detailed examination of its phenotype was not possible. However, we hypothesised that if deletion of *rrp1*^+^ is toxic in the *nmtP3-GFP-rad51* strain, *rrp1*^+^ overexpression should be beneficial. Indeed, we found that overexpression of *rrp1*^+^, but not *rrp2*^+^, rescued the growth defect and viability loss induced by Rad51 overproduction (Figure 1D,E). We confirmed that addition of the GFP tag to Rad51 was not responsible for toxicity of Rad51 overproduction or for the role of Rrp1 in alleviating this phenotype (Figure S1A). This allowed us to use a synthetic dosage lethality approach with the *nmtP3-GFP-rad51* strain to assess the effect of Rrp1 on Rad51 activity.

### The ATPase activity of Rrp1 is required to counter the genotoxicity associated with Rad51 overproduction

Rrp1 shares a complex domain structure with Uls1 (Dziadkowiec et al., 2009), so Walker B (*rrp1-DAEA*) and RING (*rrp1-CS*) Rrp1 mutant variants were employed in order to examine the importance of the respective DNA translocase and ubiquitin ligase Rrp1 activities for counteracting *rad51*^+^ overexpression-induced toxicity (Figure 2A). While the presence of a functional Rrp1 ATPase domain was required for restoration of normal growth to the *rad51*^*+*^ over-expressing strain, the RING domain appears to be dispensable (Figure 2B).

**Figure 2.**
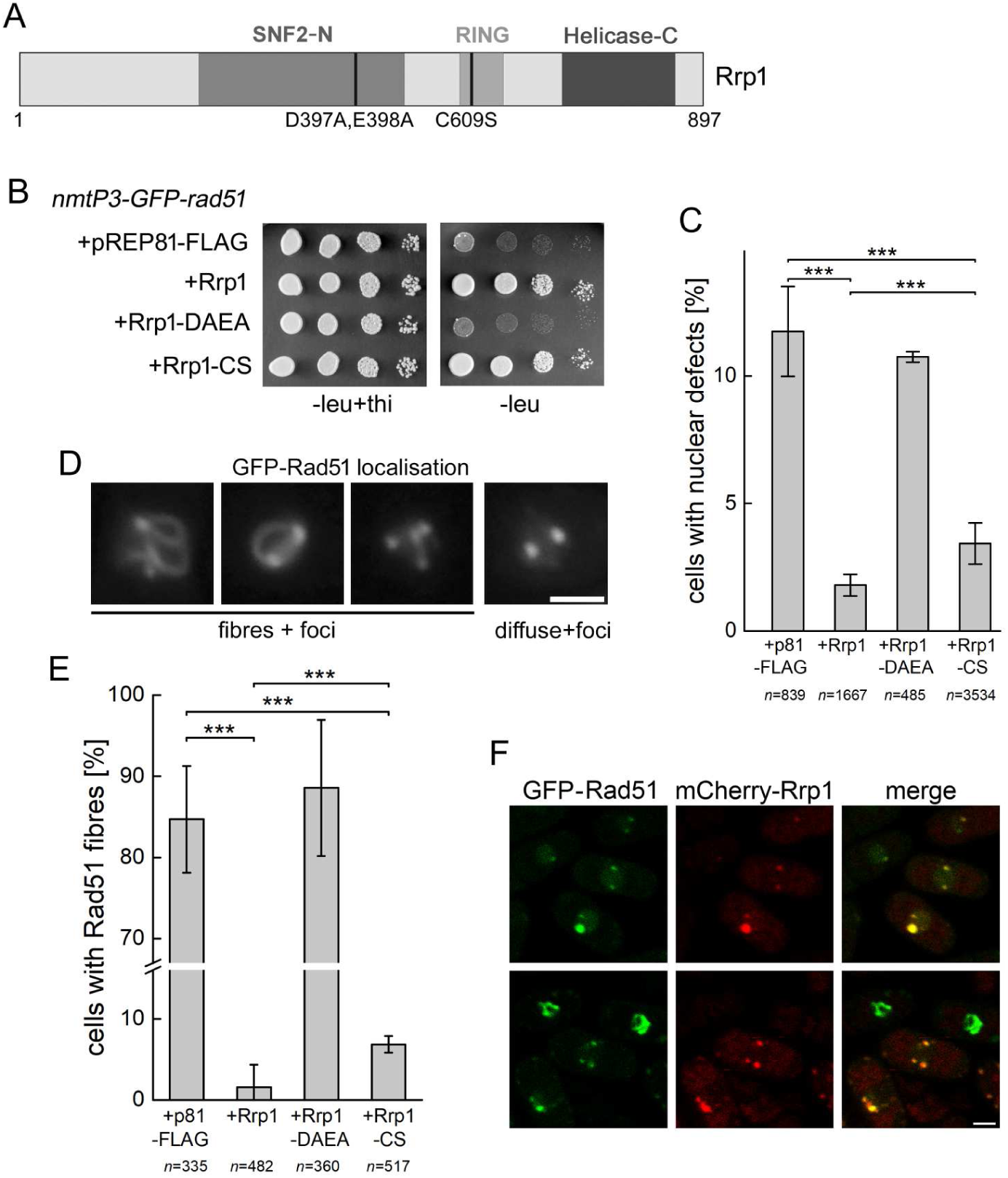
Rrp1 ATPase activity is required to suppress phenotypes caused by *rad51*^+^ overexpression. (A) Mutations abolishing Rrp1 putative SWI2/SNF2 DNA translocase (DAEA) and ubiquitin ligase (CS) activity are shown. (B) Functional Rrp1 translocase, but not RING domain, is required for suppression of the growth defect in *rad51*^+^ over-expressing cells. Cells were transformed with plasmids harbouring genes for wild type or mutated versions of respective proteins and the effect of their overexpression on the growth of cells overproducing Rad51 (-leu) or not (-leu+thi) was assessed by spot test analysis. (C) Accumulation of mitotic aberrations in *rad51*^+^ over-expressing cells is prevented by *rrp1*+ overexpression and depends on the functional translocase, but not the RING domain. Cells with unequally segregated genetic material (cut and non-disjunction) were observed by DAPI staining of the nuclei of transformants grown for 48 h in media lacking thiamine. n = total number of cells counted for 3 independent transformants of vector, *rrp1*^+^, *rrp1-DAEA or rrp1-CS*. The error bars represent the standard deviation about the mean values. The Z-test for two population proportions was used to calculate the *z*-statistic and corresponding *p*-values (*** *p* ≤ 0.001). (D) Two patterns of GFP-Rad51 localisation are shown: long fibres connecting discrete foci, and diffuse staining with foci. Scale bar represents 2 µm. (E, F) In cells overexpressing both *rrp1*^+^ and *rad51*^+^, long GFP-Rad51 fibres are no longer observed. (E) This effect is dependent on functional ATPase domain, while the RING domain may play a relatively minor role. n = total number of cells counted for 3 independent transformants of vector, *rrp1*^+^, *rrp1-DAEA or rrp1-CS* grown for 48 h in media lacking thiamine. The error bars represent the standard deviation about the mean values. The Z-test for two population proportions was used to calculate the *z*-statistic and corresponding *p*-values (*** p-value ≤ 0.001). (F) In cells overproducing both proteins, most GFP-Rad51 foci co-localise with mCherry-Rrp1 foci. Scale bar represents 2 µm.

It has previously been shown that *rad51*^+^ overexpression results in mitotic aberrations revealed by the accumulation of cells with nuclei exhibiting the cut (*cell untimely torn*) phenotype (Kim et al., 2001). We demonstrated that overproduction of Rrp1, but not the Rrp1-DAEA mutant, was able to suppress the appearance of nuclear defects that occur in the *rad51*^*+*^ overexpressing *nmtP3-GFP-rad51* strain, such as cut and non-disjunction (Figure 2C). Interestingly, the rescue induced by Rrp1-CS overproduction was less pronounced, with more cells undergoing aberrant mitosis (Figure 2C). This suggests that Rrp1 ubiquitin ligase activity might also have a role in the protein’s functional interaction with Rad51.

### Rrp1 overproduction prevents the accumulation of aberrant Rad51 staining structures

Rad51 forms only few spontaneous foci in unchallenged cells (Bernstein et al., 2011; Lorenz et al., 2009; Ouyang et al., 2009), but when overproduced, it accumulates on chromatin in human cells, forming long fibres (Mason et al., 2015; Ronneberger et al., 2008). We examined *GFP*-*rad51*^+^ over-expressing cells using fluorescence microscopy and observed that most of them contained extensive Rad51 fibres in their nuclei, often connecting several bright foci (Figure 2D). These structures virtually disappeared when Rrp1 was simultaneously overproduced, and GFP-Rad51 staining changed to diffuse with 1-3 foci (Figure 2D,E). Interestingly, about 90% of these remaining GFP-Rad51 foci co-localised with foci for Rrp1-mCherry, and GFP-Rad51 fibres were visible only in cells lacking Rrp1 signal (Figure 2F). We have previously shown that Rrp1 is enriched at centromeres and >40% of the foci it forms when overproduced are perinuclear and co-localise with Swi6 foci (Barg-Wojas et al., 2020). It is thus probable that at least some of the GFP-Rad51 foci we observe represent Rad51 bound to centromeres.

Cells overproducing Rrp1-DAEA, an ATPase deficient mutant, also contained Rad51 fibres in their nuclei (Figure 2E). This clearly demonstrates that Rrp1 translocase activity is critical to prevent excessive Rad51 accumulation on chromatin. Interestingly however, even though overproduction of the Rrp1-CS mutant did supress the toxicity of *rad51*^+^ overexpression (Figure 2B), we were nevertheless able to detect a small, but statistically significant, increase in the number of nuclei with Rad51 fibres in these cells (Figure 2E). This suggests that the putative Rrp1 ubiquitin ligase activity might also be involved in regulating Rad51 binding to DNA.

### The role of Rrp1 in regulating Rad51 function is independent from Rrp2

Overproduction of Rrp2 in a *rad51*^*+*^ overexpressing strain was unable to suppress viability loss (Figure 1D,E) and the chromosome segregation defect (Figure S1B), and did not prevent the accumulation of Rad51 fibres on chromatin (Figure S1C). Moreover, rescue of the *rad51*^*+*^ overexpression-induced growth defect by overproduction of Rrp1 was not affected by the presence of Rrp2 (Figure S1D) or the recombination auxiliary factor complex Swi5-Sfr1 (Figure S1E). This suggests that the observed role of Rrp1 in regulating Rad51 may be distinct from the previously described mutually dependent role of Rrp1 and Rrp2 in the Swi5-Sfr1 sub-pathway of HR (Dziadkowiec et al., 2009, 2013).

### Purified Rrp1 binds to DNA and has a DNA-dependent ATPase activity

In order to gain mechanistic insight into the function of Rrp1, recombinant Rrp1-FLAG was purified to near-homogeneity following overexpression in *Escherichia coli* (Figure S2A). Since Rrp1 is predicted to possess ATPase activity (Dziadkowiec et al. 2009), we first examined if the purified protein could indeed hydrolyse ATP. In the absence of DNA, no ATPase activity was detected, while robust ATP hydrolysis was observed in the presence of either ssDNA or dsDNA (Figure S2B). Based on these data, Rrp1 was estimated to have a turnover number *(k*_*cat*_) of 424 min^-1^ (± 25.1) in the presence of ssDNA and 529 min^-1^ (± 37.6) in the presence of dsDNA. These results indicate that Rrp1 possesses a robust ATPase activity that is strictly dependent on DNA, and that dsDNA stimulates ATP hydrolysis by Rrp1 more efficiently than ssDNA,

The observed dependency on DNA for ATP hydrolysis suggested that Rrp1 is capable of binding DNA. This was tested by electrophoretic mobility-shift assays (EMSA). At concentrations as low as 0.05 µM (Rrp1:nucleotide ratio of 1:100), all ssDNA was shifted in an ATP-independent manner by Rrp1, and this shift was enhanced at higher concentrations of protein (Figure S2C). A lesser shift was observed with dsDNA, with some unbound DNA remaining even at 0.30 µM Rrp1 (Rrp1:basepair ratio of 1:8.33), although ATP was also dispensable for this binding (Figure S2D). Some signal was observed in the wells, particularly for dsDNA, suggesting that Rrp1 may form aggregates consisting of protein-DNA networks that are too large to enter the gel. These results indicate that Rrp1 binds both ssDNA and dsDNA, with an apparently higher affinity for ssDNA, in an ATP-independent manner.

### Rrp1 physically interacts with Rad51

The genetic interactions observed between Rad51 and Rrp1 raised the possibility that the two proteins may physically interact. To investigate this possibility, we first employed the yeast two-hybrid system (Y2H). For Rad51, two constructs were used: a short N-terminal fragment (Rad51-N), and a long C-terminal fragment containing the Walker A and B motifs (Rad51-C) (Figure 3A). We observed a robust growth of transformants containing genes for Rrp1 and Rad51-C on high stringency SD DO-4 plates, indicating that the site of putative Rrp1 binding lies within the Rad51 region containing Walker A and B motifs (Figure 3B). In order to map the corresponding region within Rrp1 that is responsible for Rad51 binding, we created a series of four truncated forms of Rrp1 (Figure 3C) and repeated the Y2H assay with Rad51-C. These experiments revealed that the fragment of Rrp1 containing its C-terminal helicase domain was sufficient for the interaction with Rad51 (Figure 3D). In agreement with our genetic data, no interaction was observed between Rad51 and Rrp2 by Y2H (Figure 3B).

**Figure 3.**
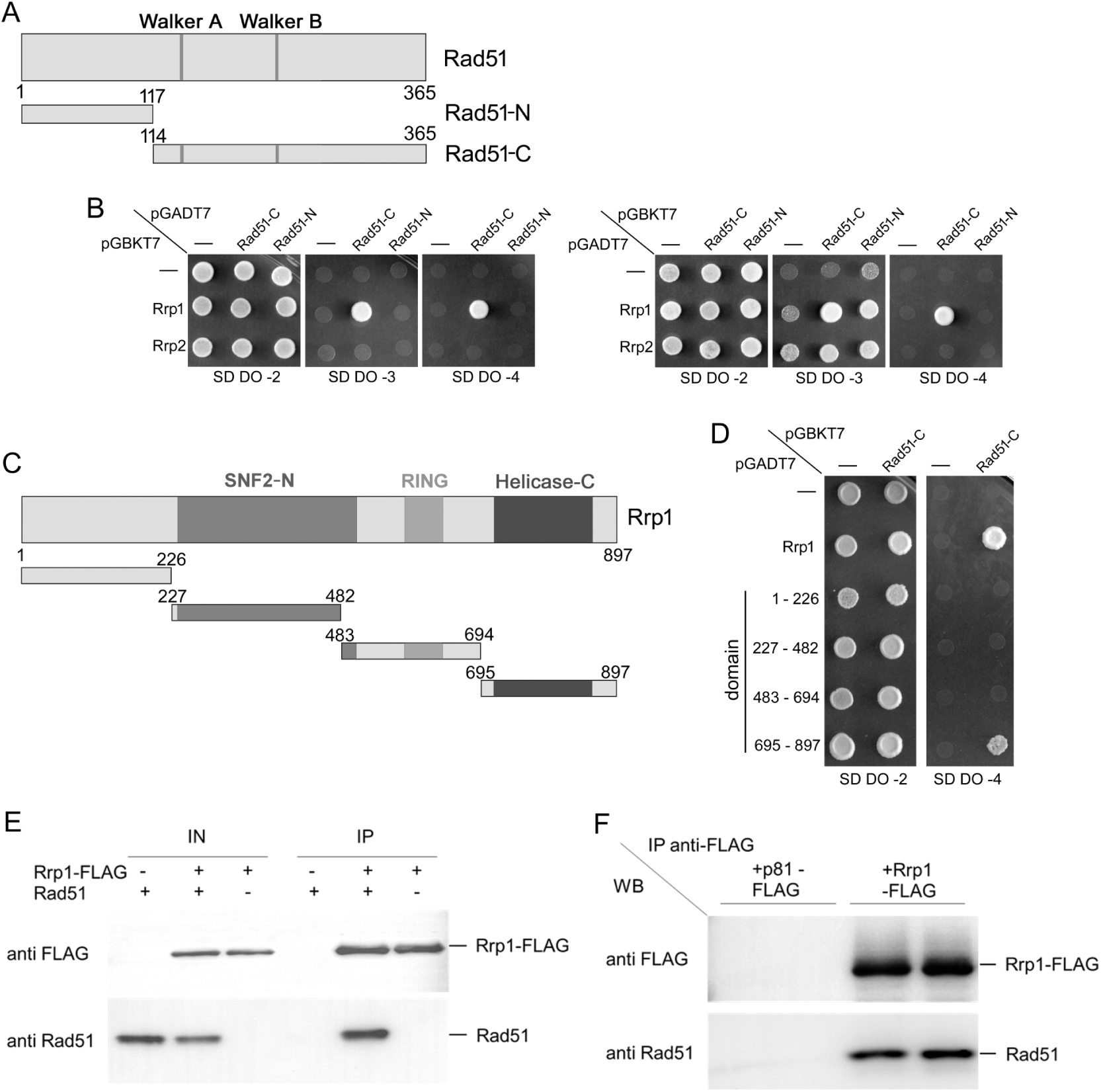
Rrp1 directly interacts with Rad51. Analysis of the Rrp1 interaction with Rad51 by yeast two-hybrid analysis. (A) Two Rad51 constructs were used: Rad51-N (N-terminal, residues 1–117) and Rad51-C (C-terminal, residues 114–365 containing the Walker A and B motifs). (B) Rad51-C (pGBKT7 and pGADT7 plasmid) may be involved in the interaction with Rrp1 (self-activation is observed in the transformant with pGADT7-Rrp1 and empty pGBKT7 vector, right panels, indicative of Rrp1 DNA binding activity). (C) Schematic representation of a series of four truncated forms of Rrp1 used to map the site of interaction with Rad51. (D) The C-terminal fragment of Rrp1 (residues 695–897) containing the C-helicase domain was found to interact with Rad51. (E) Rad51 immunoprecipitated with Rrp1-FLAG *in vitro*. Purified Rad51 and purified Rrp1-FLAG were mixed and incubated with anti-FLAG M2 agarose. Proteins were eluted from the beads with 3xFLAG peptide and separated by SDS-PAGE, then analysed by Western with anti-FLAG antibody and anti-Rad51 antiserum. -, protein was omitted and the equivalent volume of protein storage buffer was added instead. +, protein was included. (F) Rad51 interacts with Rrp1-FLAG *in vivo.* Protein extracts prepared from cells over-expressing *rrp1-FLAG* (from pREP81-FLAG plasmid) or transformed with empty vector as a control, were incubated with anti-FLAG M2 agarose. Proteins were then eluted with 3xFLAG peptide, separated by SDS-PAGE, and analysed by Western with anti-FLAG antibody and anti-Rad51 antiserum.

To validate these Y2H results and rule-out the possibility that the observed Rrp1-Rad51 interaction involved an intermediary molecule, purified Rad51 and purified Rrp1-FLAG were mixed together and subjected to immunoprecipitation with anti-FLAG M2 agarose. Importantly, Rad51 was seen to co-immunoprecipitate with Rrp1-FLAG (Figure 3E), thus confirming the existence of a direct interaction between these two proteins. Furthermore, we were able to demonstrate that Rad51-Rrp1 complex is formed *in vivo* in *S. pombe* cells by immunoprecipitating Rad51 with Rrp1-FLAG from native protein extracts (Figure 3F).

Some self-activation was observed on low stringency SD DO-3 plates when Rrp1 was fused to the transcription activation domain (pGADT7 plasmid; Figure 3B). Given the results presented here indicating that purified Rrp1 binds both ssDNA and dsDNA (Figure S2C,D), and previous microscopy and chromatin immunoprecipitation data implicating Rrp1 in DNA binding (Barg-Wojas et al., 2020), this self-activation is likely due to the ability of Rrp1 to associate with DNA.

### Rrp1 dissociates Rad51 from dsDNA *in vitro*

SWI2/SNF2-related translocases have been proposed to remove Rad51 from dsDNA in heteroduplex DNA and dead-end non-productive complexes, both in yeast and human cells (Mason et al., 2015; Shah et al., 2010). The existence of a physical interaction between the Rrp1 translocase and Rad51, combined with the ability of Rrp1 to suppress the association of Rad51 with bulk chromatin (Figure 2E), prompted us to examine whether Rrp1 can directly counteract erroneous Rad51 binding to dsDNA. To test this, we exploited the fact that the binding of purified Rrp1 to dsDNA results in a distinctive EMSA pattern where most of the dsDNA signal is retained in the well (Figure S2D). This pattern is easily distinguishable from the binding of Rad51 to DNA, which produces a smear at low concentrations of protein (0.5 µM) and a discrete band at higher concentrations (1.5 or 3 µM; Figure 4A). dsDNA was first coated with different concentrations of Rad51 and then challenged with substoichiometric amounts of Rrp1. Protein-DNA complexes were then resolved by gel agarose electrophoresis. Compared to the condition where Rrp1 was omitted, the bands for Rad51-bound DNA became fainter when 0.1 µM of Rrp1 was included. Moreover, the inclusion of 0.3 µM of Rrp1 led to a drastic decline in the signal of Rad51-dsDNA bands, even when the dsDNA was precoated with five-fold more Rad51 molecules, and signal in the well became apparent. These results suggest that sub stoichiometric amounts of Rrp1 effectively outcompete Rad51 for binding to dsDNA. Since Rrp1 can bind *in vitro* to both dsDNA (Figure S2D) and Rad51 (Figure 3E), an alternative explanation for these results is that, rather than displace Rad51 from dsDNA, Rrp1 bound to Rad51-dsDNA complexes, leading to the formation of higher-order complexes that were unable to enter the gel.

**Figure 4.**
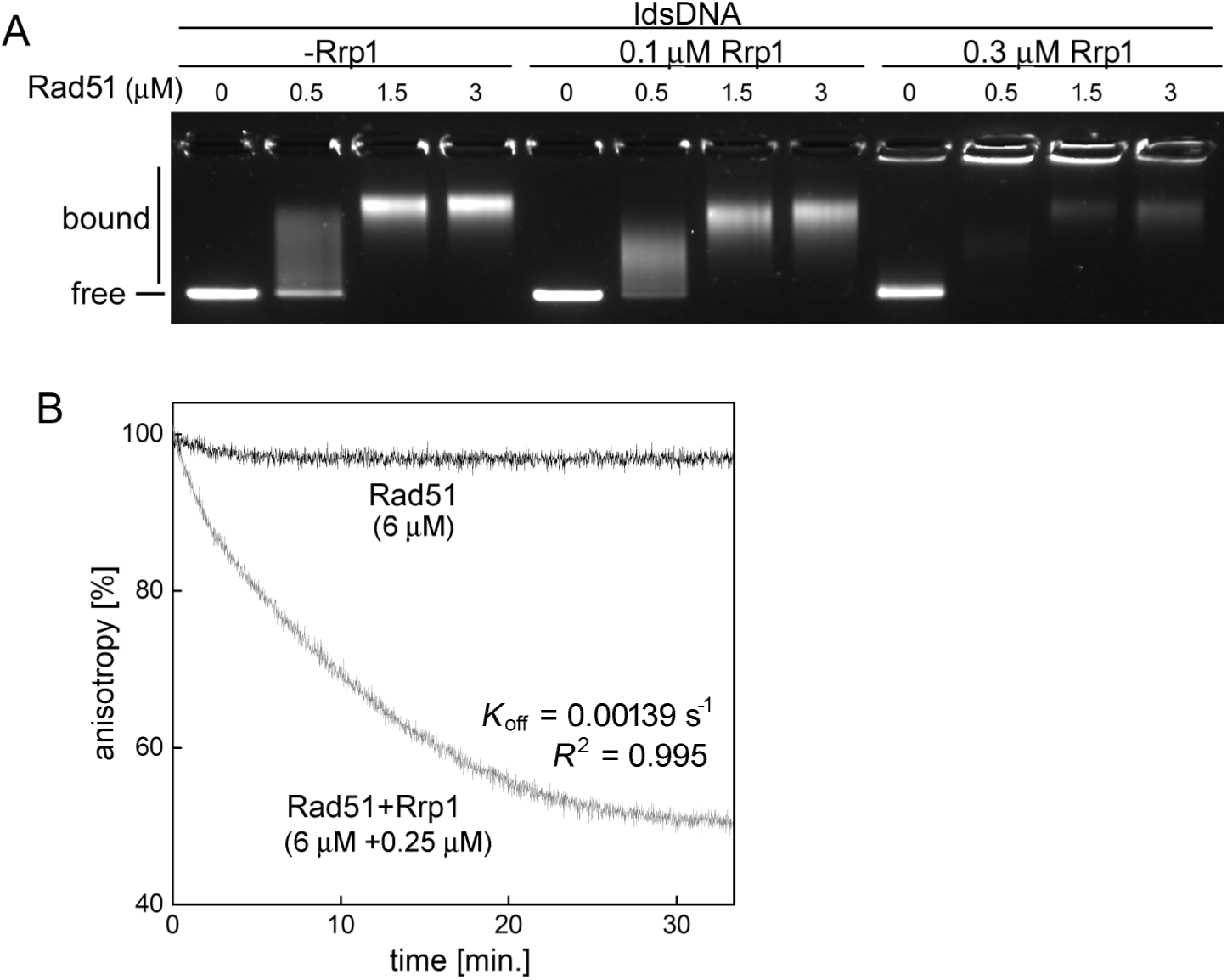
Rrp1 can disassemble Rad51-dsDNA complexes. (A) Rrp1 outcompetes Rad51 for binding to dsDNA as demonstrated by electrophoretic mobility shift assay (EMSA). Increasing amounts of Rad51 were pre-incubated with linear double-stranded DNA (ldsDNA, PhiX 174 RF I linearized with ApaLI) before addition of the indicated concentration of Rrp1. Mixtures were resolved on an agarose gel and stained with SYBR-gold. (B) Rad51-dsDNA filaments disassemble following addition of Rrp1, as demonstrated by the reduction in anisotropy of fluorescently-labelled dsDNA. Rad51 (6 µM) was incubated with a dsDNA oligonucleotide (3 µM nucleotide concentration) labelled with the TAMRA fluorophore; the resultant high anisotropy value confirms filament formation. Unlabelled heterologous scavenger DNA was then added, followed by a sub stoichiometric amount of Rrp1 (0.25 µM) or the equivalent volume of protein storage buffer, and fluorescence anisotropy was monitored for the indicated time. The decline in anisotropy observed in the reaction containing Rrp1 indicates that Rad51-dsDNA complexes are disassembled.

To more directly test whether Rrp1 could disrupt Rad51-dsDNA complexes, we employed a fluorescence anisotropy assay. Rad51-dsDNA complexes were assembled on fluorescently-labelled oligonucleotides, then Rrp1 was added and fluorescence anisotropy was monitored in real-time. When protein stock buffer was added instead of Rrp1, Rad51-dsDNA complexes remained highly stable for the duration of the experiment, with anisotropy remaining at ∼98% even after ∼30 min. By contrast, the addition of Rrp1 led to a drastic decline in anisotropy, with a value of ∼60% observed after ∼15 min (Figure 4B). This decline followed first-order decay kinetics (*R*^*2*^ = 0.995), consistent with the notion that it represents the dissociation of Rad51-dsDNA complexes, with a dissociation rate constant (*k*_*off*_) equal to 0.00139 s^-1^.

We also examined the effect of Rrp1 on Rad51-ssDNA complexes. Although a slight decrease in the intensity of Rad51-ssDNA bands was observed by EMSA (Figure S3A), this was negligible when compared with the effect of Rrp1 on Rad51-dsDNA complexes (Figure 4A). Consistently, relative anisotropy increased upon addition of Rrp1 to Rad51-ssDNA complexes in a concentration-dependent manner (Figure S3B), suggesting that rather than dissembling Rad51-dsDNA complexes, Rrp1 was binding to them.

Collectively, these results indicate that Rrp1 is capable of rapidly disassembling Rad51-dsDNA complexes. Rrp1 may also bind to Rad51-ssDNA complexes to modulate their activity but further studies are needed to conclusively test this hypothesis.

### Rrp1 has an E3 ubiquitin ligase activity with Rad51 as a substrate

In addition to ATPase domain, Rrp1 also has a Zinc finger RING-type domain characteristic of E3 ubiquitin ligases (Pombase, https://www.pombase.org/). Together with its *S. cerevisiae* orthologue Uls1, Rrp1 belongs to the RING-domain-containing Rad5/16-like group of SWI2/SNF2 translocases, distinct from the RAD54 family (Flaus and Owen-Hughes, 2011; Prasad and Ekwall, 2013). Interestingly, the pull-down of His_6_-tagged ubiquitin revealed an accumulation of ubiquitylated proteins in cells overproducing Rrp1 (Figure S4A). Moreover, when these precipitates were probed with an anti-Rad51 antibody, substantial high molecular weight species were observed (Figure S4B). We therefore hypothesized that Rrp1 may have an E3 ligase activity with Rad51 as a substrate. In order to directly test this possibility, *in vitro* Rad51 ubiquitylation assays were performed using ubiquitin, Uba1 (E1) and Ubc4 (E2) enzymes purified from *E. coli*, and purified Rrp1-FLAG as the sole E3 ubiquitin ligase enzyme. Reaction products were separated by SDS-PAGE and subjected to western blot analysis. Multiple high molecular weight protein bands were observed only when all assay components were included in the reaction (Figure 5A). Strikingly, a similar banding pattern was detected with both anti-Rad51 and anti-Ubiquitin antibodies, indicating that these bands represent ubiquitylated forms of Rad51. Consistent with this notion, the unmodified Rad51 band decreased in intensity only in the assay with all reaction components (Figure 5A). To further validate these findings, we repeated this ubiquitylation assay with either wild-type Rrp1 protein or the Rrp1-CS variant, where the Rrp1 RING domain was inactivated, as the sole E3 ligase. When Rrp1-CS was employed, the characteristic protein ladder was not formed (Figure 5B), indicating that Rrp1 E3 ubiquitin ligase activity was responsible for Rad51 ubiquitylation. Interestingly, when the membrane was probed with an anti-FLAG antibody to detect Rrp1 protein, several high molecular weight protein bands greater in size than Rrp1 were observed in reactions containing wild-type Rrp1 but not Rrp1-CS (Figure 5B, bottom panel), indicating that Rrp1 is capable of auto-ubiquitylation. Consistent with this notion, the intensity of unmodified Rrp1 band was decreased in these reactions. Interestingly, when the Rad51 ubiquitylation assay was performed in the presence of ssDNA or dsDNA, the intensity of bands corresponding to ubiquitylated Rad51 decreased (Figure 5C), with the mono-ubiquitylated form of Rad51 showing a reduction of >50%. This suggests that Rrp1 might preferably ubiquitylate free Rad51 rather than Rad51 bound to either ssDNA or dsDNA.

**Figure 5.**
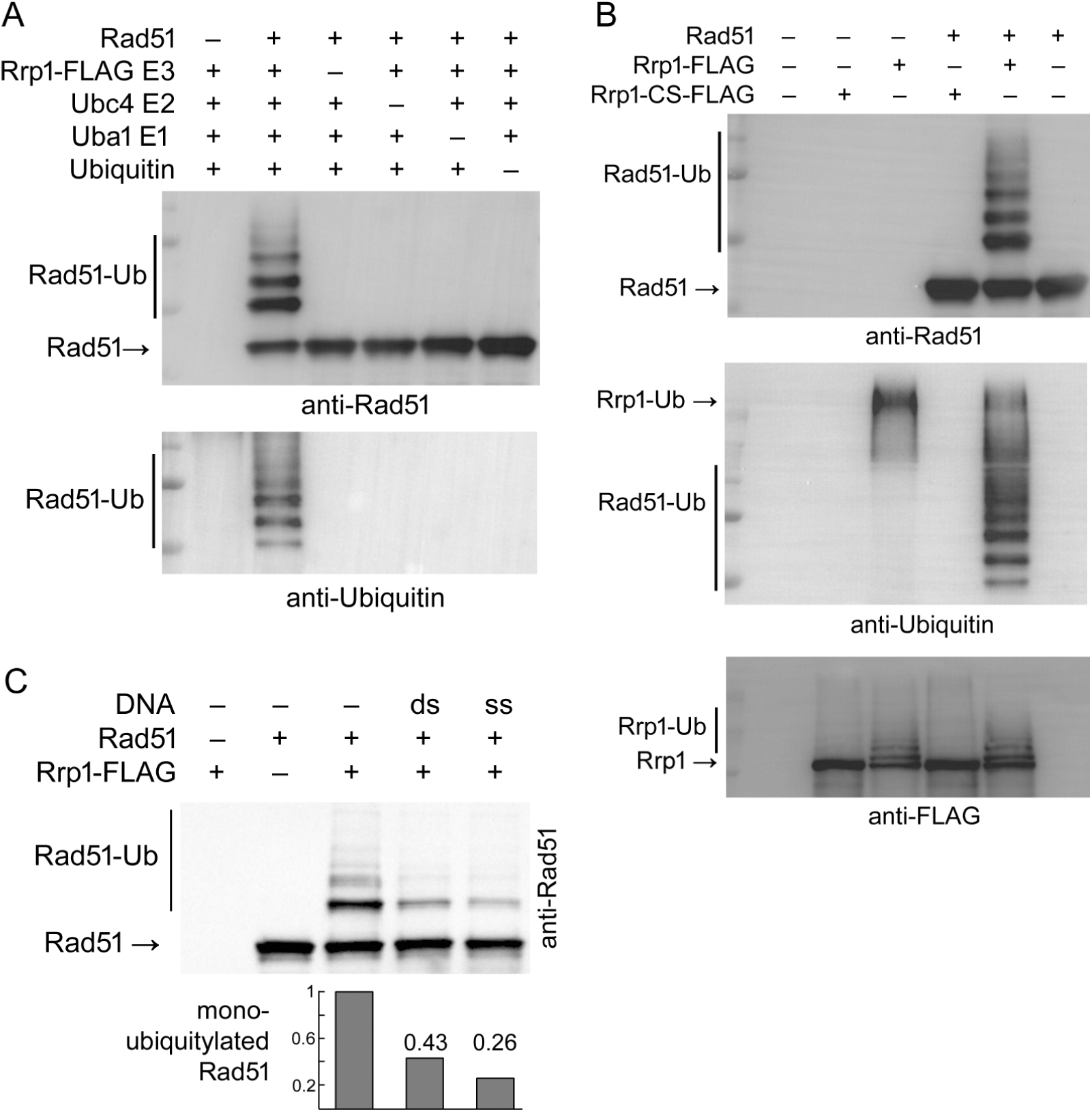
Rrp1 is an E3 ubiquitin ligase with Rad51 as a substrate. (A) The indicated reaction components were included (+) or omitted (-) for *in vitro* ubiquitylation assays. After the reaction, the reaction mixture was analysed by western blotting with anti-Rad51 antiserum and anti-Ubiquitin antibodies, and multiple bands indicative of Rad51 ubiquitylation are shown. (B) *In vitro* ubiquitylation assay containing all components as in (A) with Rrp1-FLAG or Rrp1-CS-FLAG as the E3 ligase. The reaction mixture was analysed by western blotting with anti-Rad51 antiserum and anti-Ubiquitin antibodies, demonstrating that Rrp1 RING domain is indispensable for Rad51 ubiquitylation. Additionally, reaction products were analysed with anti-FLAG antibodies (lowest panel), revealing auto-ubiquitylation of Rrp1. (C) Ubiquitylation of Rad51 by Rrp1 is less efficient in the presence of DNA. *In vitro* ubiquitylation assay containing all components as in (A) with Rad51 pre-incubated with 4 µM of ssDNA (PhiX 174 virion) or dsDNA (PhiX 174 RF I linearized with ApaLI). The reaction mixture was analysed by western blotting with anti-Rad51 antiserum and anti-Ubiquitin antibodies. The intensity ratios of mono-ubiquitylated to non-ubiquitylated Rad51 bands normalised to the sample without DNA are shown.

We thus identify Rrp1 as a translocase that can remove Rad51 from dsDNA, and a ubiquitin ligase that has Rad51 as its substrate, and propose that these Rrp1 activities are important for its role in regulating Rad51.

## Discussion

Previous studies have shown that the presence of the RAD54 family of SWI2/SNF2 DNA translocases, RAD54L and RAD54B in human (Mason et al., 2015), and Rdh54, Rad54 in *S. cerevisiae* (Shah et al., 2010), is necessary to counteract the genotoxic effects of Rad51 overproduction. Another budding yeast protein, Uls1, belonging to the RING-domain-containing Rad5/16-like group of SWI2/SNF2 DNA translocases has also been shown to participate in modulating Rad51 activity. Here we examined the interaction of *S. pombe* Rrp1, an Uls1 orthologue, with Rad51 and found that toxicity of *rad51*^+^ overexpression was increased in an *rrp1*Δ strain (Figure 1A-C). This suggested that Rrp1 has a prominent function in counteracting the toxicity of Rad51 overproduction, in contrast to the requirement for Uls1 in *S. cerevisiae*, which only becomes apparent in the absence of Rdh54 (Shah et al., 2010).

The additional viability loss conferred by deletion of *rrp1*^+^ in Rad51 overproducing cells precluded detailed examination of this mutant. However, we reasoned that concomitant *rrp1*^+^ overexpression should have the opposite effect and allow investigation of the Rrp1-Rad51 interaction. Indeed, *rrp1*^+^ overexpression rescued the growth defect and viability loss induced by Rad51 overproduction in a manner that was dependent on the presence of a functional Rrp1 ATPase domain (Figures 1D,E and 2B,C). Apart from being a DNA translocase, Uls1 was proposed to have a SUMO targeted ubiquitin ligase activity (Uzunova et al., 2007) but this role in Rad51 regulation was not examined. Rrp1 also has a Zinc finger RING-type domain characteristic of E3 ubiquitin ligases, but we found it was not crucial for rescue of the Rad51-overproduction induced growth defect.

It has been shown that Rad51 overproduction leads to chromosome segregation defects, such as chromosome bridges and micronuclei formation in mammals (Mason et al., 2015) and accumulation of cells with cut nuclei in *S. pombe* (Kim et al., 2001). We demonstrated that Rrp1 overproduction in a *rad51*^*+*^ overexpressing strain resulted in a decrease in the number of cells with cut and non-disjunction nuclear defects (Figure 2C). This was accompanied by the disappearance of extensive Rad51 fibres in the nucleus (Figure 2D,E), consistent with previous reports indicating that excessive Rad51 binding to non-damaged dsDNA may be the underlying cause of genome instability and viability loss caused by Rad51 overproduction (Mason et al., 2015; Shah et al., 2010). Rrp1-mediated rescue of these phenotypes, similarly to the viability restoration described above, was dependent on the presence of a functional Rrp1 ATPase domain, suggesting that Rrp1 modulates Rad51 through its translocase activity. Interestingly however, we found that the Rrp1-CS mutant, with an inactivated RING domain, was slightly less proficient in counteracting inappropriate Rad51 accumulation on chromatin and the appearance of aberrant DNA segregation events. This suggested that Rrp1 ubiquitin ligase activity may play a role in Rad51 regulation, even though it was apparently not important for suppression of the Rad51 overproduction-induced growth defect.

We purified Rrp1 in order to characterise its biochemical activities and obtain mechanistic insight into its function. We found that purified Rrp1 binds to both ssDNA and dsDNA independently of ATP, and has a DNA-dependent ATPase activity (Figure S2). Importantly, we demonstrated that Rrp1 and Rad51 directly interact (Figure 3) and when overproduced, they co-localise with each other in discrete perinuclear foci (Figure 2F). This supports the conclusion of our phenotypic analyses presented above, that Rad51 may be the direct target of Rrp1 activity. Indeed, we showed that Rrp1 can efficiently dissociate Rad51 from dsDNA *in vitro* (Figure 4). Thus, our data establish Rrp1 as a translocase that can counteract excessive Rad51 binding to chromatin. Such regulation by RAD54-like DNA translocases has been shown to prevent DNA segregation defects, genome instability and viability loss in mammalian cells (Mason et al., 2015), and we infer it is also the basis of Rrp1-mediated rescue of Rad51 overproduction-induced genotoxicity that we observe in *S. pombe*. Such activity has not been directly demonstrated before for Uls1 or any other protein belonging to the Rad5/16-like group of SWI2/SNF2 translocases.

The physical interaction between Rad51 and Rrp1 involves sites within the C-terminus of each protein (Figure 3B,D). Recently, the Rad51 interaction region of Sfr1, a component of the Swi5-Sfr1 HR auxiliary factor complex, was shown to be an intrinsically disordered domain (Argunhan et al., 2020). Residues 695-897 of Rrp1 involved in Rad51 binding are also predicted to include a disordered region. Thus, the intrinsically disordered structure may provide a general site for the interaction of auxiliary proteins with Rad51, but precise mapping of the interaction sites of Rrp1 will be required before direct parallels with other proteins can be drawn.

In addition to the translocase function of Rrp1, we found that the Rrp1 RING domain may also be involved in regulating Rad51 binding to DNA. We thus performed *in vitro* Rad51 ubiquitylation assays with purified Rrp1 or the Rrp1-CS RING mutant as the sole ubiquitin ligase E3 enzyme and showed that Rrp1 is able to poly-ubiquitylate Rad51 in a manner dependent on the presence of a functional Rrp1 RING domain (Figure 5). Moreover, ubiquitylated proteins accumulated in *S. pombe* cells overproducing Rrp1, and we identified Rad51 as one of these proteins (Figure S4). Taken together, this demonstrates that Rrp1 has an E3 ubiquitin ligase activity and Rad51 is one of its substrates.

This raises the interesting question of whether the Rrp1 translocase and ubiquitin ligase activities cooperate in counteracting excessive Rad51 binding to chromatin. A functional RING domain is mostly dispensable for Rrp1 mediated rescue of the Rad51 overproduction-induced growth defect, so we propose that Rad51 ubiquitylation by Rrp1 is not an absolute prerequisite for its removal from DNA by Rrp1’s translocase activity. However, the number of cells with Rad51 fibres and segregation defects increased in *rad51*^*+*^*-* overexpressing strain simultaneously overproducing Rrp1-CS compared to wild-type Rrp1.

This implies that when its ubiquitin ligase is inactivated, Rrp1 ability to prevent Rad51 association with DNA is compromised, albeit not to the extent that would affect cell viability. Interestingly, we found that addition of DNA decreased the efficiency of Rad51 ubiquitylation by Rrp1 *in vitro*, suggesting that Rrp1 may ubiquitylate free Rad51 more efficiently than Rad51 bound to either ssDNA or dsDNA. Rad51 ubiquitylation has previously been shown to compromise its ability to bind DNA (Chu et al., 2015; Inano et al., 2017). We thus hypothesise that, after the translocase activity of Rrp1 mediates Rad51 removal from chromatin, the ubiquitin ligase activity of Rrp1 could prevent the re-association of Rad51 with dsDNA. A similar model has been proposed previously for FBH1(Chu et al., 2015), a UvrD family helicase able to disrupt Rad51 nucleofilaments on ssDNA and ubiquitylate Rad51 both in human (Chu et al., 2015; Simandlova et al., 2013) and *S. pombe* cells (Tsutsui et al., 2014). This suggests that analogous strategies may be used by enzymes regulating Rad51 binding to ssDNA as well as to dsDNA.

Rad51 is overexpressed in several types of human cancers (Maacke et al., 2000a, 2000b) and multiple cancer cell lines (Hansen et al., 2003; Raderschall et al., 2002), and contributes to their increased survival after DSB induction. Increased levels of Rad51 may compensate for deficiencies in other DNA repair pathways in cancer cells and is often associated with poor patient survival prognosis (Brown and Holt, 2009; Tennstedt et al., 2013). Since the role of ubiquitylation has not been addressed in previous studies on Rad51 dysregulation (Mason et al., 2015; Shah et al., 2010), our work may have implications for patient treatment.

Although eukaryotes possess multiple SNF2 translocases, the division of labour between them is poorly understand, and even less is known about how their activities are coordinated (Poole and Cortez, 2017; Rickman and Smogorzewska, 2019). In this context, it is worth noting that Rad54 has been shown not to be required for the replication fork protection function of Rad51 (Schlacher et al., 2011; Spies et al., 2016). Budding yeast Rad5 and human SHPRH and HLTF have been shown thus far to ubiquitylate PCNA (Unk et al., 2010). Recently, SHPRH has been identified as a nucleosome E3-ubiquitin-ligase (Brühl et al., 2019), and we have shown that Rrp1 and Rrp2 are involved in modulation of nucleosome dynamics important for centromere, and thus genome, stability (Barg-Wojas et al., 2020). Our present work further extends the possible range of functions performed by SNF2 enzymes and may contribute to better understanding of their role in modulating specific subsets of activities at perturbed replication forks.

## Acknowledgments

We thank Sarah Lambert for encouraging comments on the manuscript.

## Author contributions

Jakub Muraszko, Formal analysis, Investigation, Methodology, Funding acquisition, Writing - original draft; Bilge Argunhan, Formal analysis, Methodology, Resources, Funding acquisition, Writing - review and editing; Kentaro Ito, Formal analysis, Methodology, Writing - review and editing; Gabriela Baranowska, Investigation, Funding acquisition; Anna Barg-Wojas, Karol Kramarz, Investigation; Yumiko Kurokawa, Methodology, Resources; Hiroshi Iwasaki, Supervision, Funding acquisition, Methodology, Writing - review and editing; Dorota Dziadkowiec, Conceptualization, Supervision, Formal analysis, Funding acquisition, Investigation, Methodology, Writing - original draft, Writing - review and editing.

## Materials and methods

### Yeast strains, plasmids and general methods

Strains and plasmids used are listed in Tables S1 and S2, respectively.

Media used for *S. pombe* growth were as described (Moreno et al., 1991). Yeast cells were grown at 28°C in complete yeast extract plus supplements (YES) medium or glutamate supplemented Edinburgh minimal medium (EMM). Where required, thiamine was added at 5 μg/mL and geneticin (ICN Biomedicals) at 100 μg/mL. Multiple mutants were obtained by genetic crossing of relevant single mutants followed either by random spore analysis or by tetrad dissection. pREP81-FLAG vector and plasmids carrying wild type and mutated forms of *rrp1*^+^ and *rrp2*^+^ as well as domains of *rrp1*+ gene used in the yeast two hybrid system were constructed using the Gibson Assembly® Cloning Kit (NEB). All primers used to amplify gene sequences by PCR are listed in Table S3. Amplified fragments were cloned into NdeI and BamHI digested pREP81 vector. After Gibson cloning, inserts were cut by NdeI and SmaI digestion and cloned into pREP42-HA, pREP42-EGFP, pREP41-mCherry or pGADT7 and pGBKT7 plasmids. *rrp1*^+^ and *rrp1-CS* mutant version were introduced into the pGEX vector(GE Healthcare) by In-Fusion® cloning (Takara Bio). All constructs were checked by sequencing.

### Spot assays

Cells were grown to mid-log phase, then serially diluted by 10-fold and 2 µL aliquots were spotted onto relevant plates (YES or EMM) that were incubated for 3-5 days in 28°C and photographed. All assays were repeated at least twice.

### Survival assay

Cells were grown for 48 hours in YES or in minimal medium with (repressed conditions) or without thiamine (overexpression) at 28°C. 500 µL aliquots were collected, serially diluted and plated onto YES plates to determine the number of viable cells. Plates were incubated for 3-5 days at 28°C. The viable cells were counted and percentage of survival for gene overexpression conditions was calculated against the repressed control.

### Yeast two-hybrid assay

Gal4-based Matchmaker Two-Hybrid System 3 (Clontech) was used according to manufacturer’s instructions. The indicated proteins were fused to the GAL4 activation domain (AD) in pGADT7 vector and the GAL4 DNA-binding domain (DBD) in pGBKT7, and expressed in the *S. cerevisiae* tester strain AH109. Transformants were selected on synthetic dextrose drop-out medium without Leu and Trp (SD DO-2), and then plated on low stringency medium without Leu, Trp and His (SD DO-3) and high stringency medium without Leu, Trp, His and Ade (SD DO-4), and incubated for 3-5 days at 28°C.

### Fluorescence microscopy

To determine the formation of Rrp1 foci and their co-localisation with Rad51 foci, appropriate transformants were grown for 24 h in EMM medium without thiamine. 1 mL of culture was harvested, washed with water and subjected to fluorescent microscopy analysis. For co-localisation experiments, data were collected under 63x magnification with the confocal microscope Leica 453 TCS SP8 (Leica Microsystems) equipped with Leica HyD SP detector, and analysed with LAS X 3.3.0. For examination of mitotic defects induced by *rad51*^+^ overexpression, samples taken from respective transformant cultures grown for 48 hours in EMM medium without thiamine were washed and fixed in 70% ethanol. After rehydration, cells were stained with 1 mg/mL 4’,6-diamidino-2-phenylindole (DAPI) and 1 mg/mL p-phenylenediamine in 50% glycerol and examined by fluorescence microscopy with Axio Imager A.2 (Carl Zeiss).

### *In vivo* co-immunoprecipitation

*In vivo* pull-down experiments were performed using strains with native levels of Rad51 and overproduction of FLAG-tagged Rrp1, which was integrated into the *ars1* region under control of the *nmt81* promotor. Cells were grown to mid-log phase using EMM minimal medium for 24 hours. 100 mL of cells were harvested and broken with glass beads in H buffer (50 mM HEPES-KOH pH 7, 50 mM KOAc, 5 mM MgOAc, 0.1% NP-40, 10% glycerol, 1 mM DTT and 1x cOmplete™ EDTA-free protease inhibitor cocktail (Roche)). Extracts were cleared by centrifugation and immunoprecipitated with ANTI-FLAG® M2 Affinity Gel (Sigma). Beads were washed and eluted using 100 µg/mL 3xFLAG peptide (Sigma). For detection, anti-FLAG (1:5000, Sigma) antibodies and anti-Rad51 (1:5000) (Haruta et al., 2006) antiserum were used.

### Purification of Rrp1-FLAG

Recombinant Rrp1 and Rrp1-CS were expressed in the Rosetta *E. coli* strain (Novagen) from the pGEX-6P plasmid (GE Healthcare). Proteins were C-terminally fused to the GST tag and N-terminally to the 3xFLAG tag; only the former was removed during the purification process. Expression was induced with 1 mM IPTG (Sigma) at 18°C overnight. Cells were collected by centrifugation, resuspended in R buffer (20 mM Tris, pH 7.5, 10% glycerol, 1mM EDTA) containing 500 mM NaCl, and disrupted by sonication. The cell lysate was then clarified by ultracentrifugation (70,000 *g*, 1 h, 2°C). The supernatant was mixed with 4B GSH sepharose (Sigma) for 3 h at 4°C. Resin-bound proteins were eluted in R buffer containing 300 mM NaCl and 40 mM glutathione (Sigma). The sample was supplemented with 0.2 µg/mL of HRV-3C protease (Sigma) to remove the GST tag and dialyzed against R buffer containing 100 mM NaCl (overnight, 4°C). The dialyzed sample was loaded onto a 1 mL HiTrap Heparin (Sigma) column. Rrp1 eluted at around 650 mM NaCl with a linear gradient of 0.1-1.0 M NaCl in R buffer. Eluted fractions were diluted 6.5-fold with R buffer and loaded onto a 1 mL ResourceQ column (GE Healthcare). Rrp1 eluted at around 300 mM NaCl with a linear gradient of 0.1-1.0 M NaCl in R buffer. Eluted fractions were diluted 3-fold with R buffer and loaded onto a HiTrap SP column (GE Healthcare). Rrp1 eluted at around 600 mM NaCl with a linear gradient of 0.1-1.0 M NaCl in R buffer. Eluted fractions were collected and dialysed against R buffer containing 200 mM NaCl. Concentration was determined using NanoDrop (ThermoFisher) with a molar extinction coefficient of 100365. For the Rrp1-CS mutant, Resource Q and HiTrap SP columns were omitted. Rad51 was purified exactly as previously described (Kurokawa et al., 2008). Uba1 (E1) and Ubc4 (E2) were purified exactly as previously described (Tsutsui et al., 2014). All proteins were free of nuclease and/or protease activities for the duration of the relevant assays.

### *In vitro* co-immunoprecipitation

*in vitro* co-immunoprecipitation assays were performed as follows. Purified Rad51 and FLAG-tagged Rrp1 (250 nM each) were incubated in IP buffer (35 mM Tris-HCl pH 7.5, 1 mM DTT, 100 mM NaCl, 3.5 mM MgCl_2_, 0.1% NP-40, 5% glycerol) for 30 min at 30°C. Proteins were immunoprecipitated using ANTI-FLAG® M2 Affinity Gel (Sigma) for 2 h at 4°C. Beads were washed three times with IP buffer, and proteins were eluted using 100 µg/mL 3x FLAG peptide (Sigma). Eluates were analysed by western blotting with anti-FLAG antibodies (1:5000, Sigma) and anti-Rad51 (1:5000),(Haruta et al., 2006) antiserum.

### Colorimetric ATPase assay

ATPase activity was measured using a commercial malachite green phosphate detection kit (BioAssay Systems) according to the manufacturer’s instructions. Purified Rrp1 (30 nM) was incubated with or without 5 µM nucleotides DNA substrates (cssDNA, Phi X174; or ldsDNA, Phi 174 RF I; both from NEB) with 1 mM ATP in ATPase buffer (25 mM Tris-HCl [pH 7.5], 1 mM DTT, 20 mM NaCl, 5 mM MgCl_2_ and 2% glycerol) at 30°C. The reaction was stopped at specific time points via addition of 2.25 M H_2_SO_4_.

### Electrophoretic mobility shift assay (EMSA)

Purified Rrp1 was incubated with ssDNA (Phi X174, NEB) or dsDNA (Phi 174 RF I, NEB; linearized with ApaLI) in E buffer (25 mM HEPES pH 7.5, 1 mM DTT, 60 mM KCl, 2 mM ATP, 3.5 mM MgCl_2_, 5% glycerol) for 15 min in 37°C. Samples were then crosslinked with 0.2% glutaraldehyde (37°C, 5 min) then run on a 0.8% agarose 1xTAE gel and stained with SYBR Gold (ThermoFisher).

### *In vitro* ubiquitylation assay

Reactions were performed in ubiquitylation buffer (25 mM Tris-HCl [pH 7.5], 1 mM DTT, 20 mM NaCl, 5 mM ATP, 5 mM MgCl_2_ and 2% glycerol) containing 5 µM *S. cerevisiae* ubiquitin (Funakoshi, U-100SC-05M), 0.2 µM His-Uba1 (E1), 2 µM His-Ubc4 (E2), 0.2 µM Rrp1-FLAG (E3) and 1 µM Rad51. This mixture was incubated at 37°C for 30 min. When the effect of DNA was examined, Rad51 was preincubated with 4 µM of ssDNA (PhiX virion) or dsDNA (PhiX RFI linearised with ApaLI) at 37°C for 20 min in ubiquitylation buffer prior to the addition of ubiquitylation reaction components. Reactions were stopped by addition of 6x SDS Laemmli buffer and proteins were resolved by 15% SDS-PAGE and analysed by western blotting with anti-ubiquitin antibodies (1:2000 Abcam P4D1) and anti-Rad51 (1:5000) (Haruta et al., 2006) antiserum.

### Rad51 removal from DNA

EMSA-based analysis was performed as described above except that Rad51 was first incubated with the DNA at 37°C for 5 min, then Rrp1 was added. Fluorescence anisotropy analysis was as follows. For the measurements with ssDNA Rad51 filaments were formed on by incubation of 0.5 µM Rad51 with 1.5 µM ssDNA (oligo-dT, 72-mer) for 5 minutes at 37°C in buffer (30 mM HEPES pH 7.5, 1 mM DTT, 50 mM NaCl, 100 mM KCl, 2 mM ATP, 8 mM PC, 8 U/ml CPK, 3.5 mM MgCl_2_, 2.5% glycerol). This mixture was transferred to a 0.2 × 1.0 cm cuvette (Hellma Analytics) at 37°C. The change in fluorescence anisotropy at 575 nm following excitation at 546 nm was measured for 60 s. After that time, competitor DNA (15 µM nucleotides PhiX 174 virion) and the indicated concentrations of Rrp1 protein were added. Data were collected using an FP-8300 spectrofluorometer (JASCO) every second for 3 minutes. For each reaction, the measurements 20 s before addition of scavenger DNA and Rrp1 were averaged, and the subsequent measurements were expressed relative to this averaged value.

For the measurements with dsDNA 6 µM of Rad51 was incubated with 3 µM (base pair concentration) of dsDNA (5’-TAMRA-AAATG AACAT AAAGT AAATA AGTAT AAGGA TAATA CAAAA TAAGT AAATG AATAA ACATA GAAAA TAAAG TA-3’ annealed to its complementary strand) for 5 minutes at 37°C in buffer (30 mM HEPES pH 7.5, 1 mM DTT, 50 mM NaCl, 100 mM KCl, 2 mM ATP, 8 mM PC, 8 U/ml CPK, 3.5 mM MgCl_2_, 2.5% glycerol). This mixture was then transferred to a 1.0 × 1.0 cm cuvette (Hellma Analytics) with a magnetic stirrer and maintained at 37°C with stirring (450rpm). The change in fluorescence anisotropy at 575 nm following excitation at 546 nm was monitored for 60 s. After that time, scavenger DNA (15 µM nucleotides PhiX 174 virion) was added, and after 20 s, 0.25 µM of Rrp1 was injected into the mixture. Data were collected using an FP-8300 spectrofluorometer (JASCO) every second for over 30 min. The dissociation rate constant (*k*_*off*_) was calculated using the following equation:

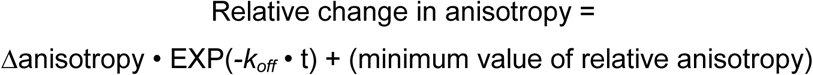

where Δanisotropy in the difference between anisotropy at time zero (average of measurements 20 s before addition of scavenger DNA) and the lowest value observed in the experiment, and Relative change in anisotropy is the change in anisotropy at time t relative to time zero.

### Statistical data analysis

For viability assays, Student’s t test was used to calculate the P-values (*** *p* ≤ 0.001, ** 0.001 < *p* ≤ 0.01, * 0.01 < *p* ≤ 0.05). To assess statistical significance of proportions of cells with aberrant mitosis and the Rad51 localisation pattern, the Z-test for two population proportions was used to calculate the *z*-statistic and corresponding *p*-values (*** *p* ≤ 0.001, ** 0.001 < *p* ≤ 0.01, * 0.01 < *p* ≤ 0.05).

## References

Achar, Y.J., Balogh, D., Neculai, D., Juhasz, S., Morocz, M., Gali, H., Dhe-Paganon, S., Venclovas, Č., and Haracska, L. (2015). Human HLTF mediates postreplication repair by its HIRAN domain-dependent replication fork remodelling. Nucleic Acids Res. 43, 10277–10291.

Akamatsu, Y., Tsutsui, Y., Morishita, T., Siddique, M.S.P., Kurokawa, Y., Ikeguchi, M., Yamao, F., Arcangioli, B., and Iwasaki, H. (2007). Fission yeast Swi5/Sfr1 and Rhp55/Rhp57 differentially regulate Rhp51-dependent recombination outcomes. EMBO J. 26, 1352–1362.

Argunhan, B., Murayama, Y., and Iwasaki, H. (2017). The differentiated and conserved roles of Swi5-Sfr1 in homologous recombination. FEBS Lett. 591, 2035–2047.

Argunhan, B., Sakakura, M., Afshar, N., Kurihara, M., Ito, K., Maki, T., Kanamaru, S., Murayama, Y., Tsubouchi, H., Takahashi, M., et al. (2020). Cooperative interactions facilitate stimulation of Rad51 by the Swi5-Sfr1 auxiliary factor complex. Elife 9.

Barg-Wojas, A., Muraszko, J., Kramarz, K., Schirmeisen, K., Baranowska, G., Carr, A.M., and Dziadkowiec, D. (2020). Schizosaccharomyces pombe DNA translocases Rrp1 and Rrp2 have distinct roles at centromeres and telomeres that ensure genome stability. J. Cell Sci. 133, jcs230193.

Bennett, B.T., and Knight, K.L. (2005). Cellular localization of human Rad51C and regulation of ubiquitin-mediated proteolysis of Rad51. J. Cell. Biochem. 96, 1095–1109.

Bernstein, K.A., Reid, R.J.D., Sunjevaric, I., Demuth, K., Burgess, R.C., and Rothstein, R. (2011). The Shu complex, which contains Rad51 paralogues, promotes DNA repair through inhibition of the Srs2 anti-recombinase. Mol. Biol. Cell 22, 1599–1607.

Bétous, R., Mason, A.C., Rambo, R.P., Bansbach, C.E., Badu-Nkansah, A., Sirbu, B.M., Eichman, B.F., and Cortez, D. (2012). SMARCAL1 catalyzes fork regression and Holliday junction migration to maintain genome stability during DNA replication. Genes Dev. 26, 151–162.

Bhat, K.P., and Cortez, D. (2018). RPA and RAD51: Fork reversal, fork protection, and genome stability. Nat. Struct. Mol. Biol. 25, 446–453.

Bhat, K.P., Krishnamoorthy, A., Dungrawala, H., Garcin, E.B., Modesti, M., and Cortez, D. (2018). RADX Modulates RAD51 Activity to Control Replication Fork Protection. Cell Rep. 24, 538–545.

Brown, E.T., and Holt, J.T. (2009). Rad51 overexpression rescues radiation resistance in BRCA2-defective cancer cells. Mol. Carcinog. 48, 105–109.

Brühl, J., Trautwein, J., Schäfer, A., Linne, U., and Bouazoune, K. (2019). The DNA repair protein SHPRH is a nucleosome-stimulated ATPase and a nucleosome-E3 ubiquitin ligase. Epigenetics and Chromatin 12, 1–16.

Bugreev, D. V, Rossi, M.J., and Mazin, A. V (2011). Cooperation of RAD51 and RAD54 in regression of a model replication fork. Nucleic Acids Res. 39, 2153–2164.

Carr, A.M., Paek, A.L., and Weinert, T. (2011). DNA replication: failures and inverted fusions. Semin. Cell Dev. Biol. 22, 866–874.

Chi, P., Kwon, Y., Seong, C., Epshtein, A., Lam, I., Sung, P., and Klein, H.L. (2006). Yeast recombination factor Rdh54 functionally interacts with the Rad51 recombinase and catalyzes Rad51 removal from DNA. J. Biol. Chem. 281, 26268–26279.

Chi, P., Kwon, Y., Visnapuu, M.-L., Lam, I., Santa Maria, S.R., Zheng, X., Epshtein, A., Greene, E.C., Sung, P., and Klein, H.L. (2011). Analyses of the yeast Rad51 recombinase A265V mutant reveal different in vivo roles of Swi2-like factors. Nucleic Acids Res. 39, 6511–6522.

Chu, W.K., Payne, M.J., Beli, P., Hanada, K., Choudhary, C., and Hickson, I.D. (2015). FBH1 influences DNA replication fork stability and homologous recombination through ubiquitylation of RAD51. Nat. Commun. 6, 5931.

Dungrawala, H., Bhat, K.P., Le Meur, R., Chazin, W.J., Ding, X., Sharan, S.K., Wessel, S.R., Sathe, A.A., Zhao, R., and Cortez, D. (2017). RADX Promotes Genome Stability and Modulates Chemosensitivity by Regulating RAD51 at Replication Forks. Mol. Cell 67, 374-386.e5.

Dziadkowiec, D., Petters, E., Dyjankiewicz, A., Karpiński, P., Garcia, V., Watson, A.T., and Carr, A.M. (2009). The role of novel genes rrp1(+) and rrp2(+) in the repair of DNA damage in Schizosaccharomyces pombe. DNA Repair (Amst). 8, 627–636.

Dziadkowiec, D., Kramarz, K., Kanik, K., Wiśniewski, P., and Carr, A.M. (2013). Involvement of Schizosaccharomyces pombe rrp1+ and rrp2 + in the Srs2- and Swi5/Sfr1-dependent pathway in response to DNA damage and replication inhibition. Nucleic Acids Res. 41, 8196–8209.

Flaus, A., and Owen-Hughes, T. (2011). Mechanisms for ATP-dependent chromatin remodelling: The means to the end. FEBS J. 278, 3579–3595.

Hansen, L.T., Lundin, C., Spang-Thomsen, M., Petersen, L.N., and Helleday, T. (2003). The role of RAD51 in etoposide (VP16) resistance in small cell lung cancer. Int. J. Cancer 105, 472–479.

Haruta, N., Kurokawa, Y., Murayama, Y., Akamatsu, Y., Unzai, S., Tsutsui, Y., and Iwasaki, H. (2006). The Swi5-Sfr1 complex stimulates Rhp51/Rad51- and Dmc1-mediated DNA strand exchange in vitro. Nat. Struct. Mol. Biol. 13, 823–830.

Hashimoto, Y., Ray Chaudhuri, A., Lopes, M., and Costanzo, V. (2010). Rad51 protects nascent DNA from Mre11-dependent degradation and promotes continuous DNA synthesis. Nat. Struct. Mol. Biol. 17, 1305–1311.

Heyer, W.-D. (2015). Regulation of recombination and genomic maintenance. Cold Spring Harb. Perspect. Med. 5, 1–24.

Inano, S., Sato, K., Katsuki, Y., Kobayashi, W., Tanaka, H., Nakajima, K., Nakada, S., Miyoshi, H., Knies, K., Takaori-Kondo, A., et al. (2017). RFWD3-Mediated Ubiquitination Promotes Timely Removal of Both RPA and RAD51 from DNA Damage Sites to Facilitate Homologous Recombination. Mol. Cell 66, 622-634.e8.

Kile, A.C., Chavez, D.A., Bacal, J., Eldirany, S., Korzhnev, D.M., Bezsonova, I., Eichman, B.F., and Cimprich, K.A. (2015). HLTF’s Ancient HIRAN Domain Binds 3’ DNA Ends to Drive Replication Fork Reversal. Mol. Cell 58, 1090–1100.

Kim, W.J., Lee, H., Park, E.J., Park, J.K., and Park, S.D. (2001). Gain- and loss-of-function of Rhp51, a Rad51 homolog in fission yeast, reveals dissimilarities in chromosome integrity. Nucleic Acids Res. 29, 1724–1732.

Kolinjivadi, A.M., Sannino, V., de Antoni, A., Técher, H., Baldi, G., and Costanzo, V. (2017). Moonlighting at replication forks – a new life for homologous recombination proteins BRCA1, BRCA2 and RAD51. FEBS Lett. 591, 1083–1100.

Kowalczykowski, S.C. (2015). An Overview of the Molecular Mechanisms of Recombinational DNA Repair. Cold Spring Harb. Perspect. Biol. 7, a016410.

Krogh, B.O., and Symington, L.S. (2004). Recombination proteins in yeast. Annu. Rev. Genet. 38, 233–271.

Kurokawa, Y., Murayama, Y., Haruta-Takahashi, N., Urabe, I., and Iwasaki, H. (2008). Reconstitution of DNA strand exchange mediated by Rhp51 recombinase and two mediators. PLoS Biol. 6, e88.

Lambert, S., Watson, A.T., Sheedy, D.M., Martin, B., and Carr, A.M. (2005). Gross chromosomal rearrangements and elevated recombination at an inducible site-specific replication fork barrier. Cell 121, 689–702.

Liu, J., and Heyer, W.-D. (2011). Who’s who in human recombination: BRCA2 and RAD52. Proc. Natl. Acad. Sci. U. S. A. 108, 441–442.

Lorenz, A., Osman, F., Folkyte, V., Sofueva, S., and Whitby, M.C. (2009). Fbh1 Limits Rad51-Dependent Recombination at Blocked Replication Forks. Mol. Cell. Biol. 29, 4742–4756.

Maacke, H., Jost, K., Opitz, S., Miska, S., Yuan, Y., Hasselbach, L., Lüttges, J., Kalthoff, H., and Stürzbecher, H.-W. (2000a). DNA repair and recombination factor Rad51 is over-expressed in human pancreatic adenocarcinoma. Oncogene 19, 2791–2795.

Maacke, H., Opitz, S., Jost, K., Hamdorf, W., Henning, W., Krüger, S., Feller, A.C., Lopens, A., Diedrich, K., Schwinger, E., et al. (2000b). Over-expression of wild-type Rad51 correlates with histological grading of invasive ductal breast cancer. Int. J. Cancer 88, 907–913.

Mason, J.M., Dusad, K., Wright, W.D., Grubb, J., Budke, B., Heyer, W.-D., Connell, P.P., Weichselbaum, R.R., and Bishop, D.K. (2015). RAD54 family translocases counter genotoxic effects of RAD51 in human tumor cells. Nucleic Acids Res. 43, 3180–3196.

McGlynn, P., and Lloyd, R.G. (2002). Recombinational repair and restart of damaged replication forks. Nat. Rev. Mol. Cell Biol. 3, 859–870.

Mijic, S., Zellweger, R., Chappidi, N., Berti, M., Jacobs, K., Mutreja, K., Ursich, S., Ray Chaudhuri, A., Nussenzweig, A., Janscak, P., et al. (2017). Replication fork reversal triggers fork degradation in BRCA2-defective cells. Nat. Commun. 8, 859.

Moreno, S., Klar, A., and Nurse, P. (1991). Molecular genetic analysis of fission yeast Schizosaccharomyces pombe. Methods Enzymol. 194, 795–823.

Ouyang, K.J., Woo, L.L., Zhu, J., Huo, D., Matunis, M.J., and Ellis, N.A. (2009). SUMO modification regulates BLM and RAD51 interaction at damaged replication forks. PLoS Biol. 7.

Pepe, A., and West, S.C. (2014). MUS81-EME2 promotes replication fork restart. Cell Rep. 7, 1048–1055.

Piwko, W., Mlejnkova, L.J., Mutreja, K., Ranjha, L., Stafa, D., Smirnov, A., Brodersen, M.M., Zellweger, R., Sturzenegger, A., Janscak, P., et al. (2016). The MMS22L–TONSL heterodimer directly promotes RAD51-dependent recombination upon replication stress. EMBO J. 35, 2584–2601.

Poole, L.A., and Cortez, D. (2017). Functions of SMARCAL1, ZRANB3, and HLTF in maintaining genome stability. Crit. Rev. Biochem. Mol. Biol. 52, 696–714.

Prakash, R., Zhang, Y., Feng, W., and Jasin, M. (2015). Homologous recombination and human health: the roles of BRCA1, BRCA2, and associated proteins. Cold Spring Harb. Perspect. Biol. 7, a016600.

Prasad, P., and Ekwall, K. (2013). A snapshot of Snf2 enzymes in fission yeast. Biochem. Soc. Trans. 41, 1640–1647.

Raderschall, E., Stout, K., Freier, S., Suckow, V., Schweiger, S., and Haaf, T. (2002). Elevated levels of Rad51 recombination protein in tumor cells. Cancer Res. 62, 219–225.

Raji, H., and Hartsuiker, E. (2006). Double-strand break repair and homologous recombination in Schizosaccharomyces pombe. Yeast 23, 963–976.

Rickman, K., and Smogorzewska, A. (2019). Advances in understanding DNA processing and protection at stalled replication forks. J. Cell Biol. 218, 1096–1107.

Ronneberger, O., Baddeley, D., Scheipl, F., Verveer, P.J., Burkhardt, H., Cremer, C., Fahrmeir, L., Cremer, T., and Joffe, B. (2008). Spatial quantitative analysis of fluorescently labeled nuclear structures: problems, methods, pitfalls. Chromosome Res. 16, 523–562.

Schlacher, K., Christ, N., Siaud, N., Egashira, A., Wu, H., and Jasin, M. (2011). Double-strand break repair-independent role for BRCA2 in blocking stalled replication fork degradation by MRE11. Cell 145, 529–542.

Shah, P.P., Zheng, X., Epshtein, A., Carey, J.N., Bishop, D.K., and Klein, H.L. (2010). Swi2/Snf2-related translocases prevent accumulation of toxic Rad51 complexes during mitotic growth. Mol. Cell 39, 862–872.

Simandlova, J., Zagelbaum, J., Payne, M.J., Chu, W.K., Shevelev, I., Hanada, K., Chatterjee, S., Reid, D.A., Liu, Y., Janscak, P., et al. (2013). FBH1 helicase disrupts RAD51 filaments in vitro and modulates homologous recombination in mammalian cells. J. Biol. Chem. 288, 34168–34180.

Solinger, J.A., Kiianitsa, K., and Heyer, W.-D. (2002). Rad54, a Swi2/Snf2-like Recombinational Repair Protein, Disassembles Rad51:dsDNA Filaments. Mol. Cell 10, 1175–1188.

Spies, J., Waizenegger, A., Barton, O., Sürder, M., Wright, W.D., Heyer, W.-D., and Löbrich, M. (2016). Nek1 Regulates Rad54 to Orchestrate Homologous Recombination and Replication Fork Stability. Mol. Cell 903–917.

Sung, P., and Klein, H.L. (2006). Mechanism of homologous recombination: mediators and helicases take on regulatory functions. Nat. Rev. Mol. Cell Biol. 7, 739–750.

Sung, P., and Robberson, D.L. (1995). DNA strand exchange mediated by a RAD51-ssDNA nucleoprotein filament with polarity opposite to that of RecA. Cell 82, 453–461.

Symington, L.S. (2002). Role of RAD52 epistasis group genes in homologous recombination and double-strand break repair. Microbiol. Mol. Biol. Rev. 66, 630–670.

Tennstedt, P., Fresow, R., Simon, R., Marx, A., Terracciano, L., Petersen, C., Sauter, G., Dikomey, E., and Borgmann, K. (2013). RAD51 overexpression is a negative prognostic marker for colorectal adenocarcinoma. Int. J. Cancer 132, 2118–2126.

Tsutsui, Y., Kurokawa, Y., Ito, K., Siddique, M.S.P., Kawano, Y., Yamao, F., and Iwasaki, H. (2014). Multiple Regulation of Rad51-Mediated Homologous Recombination by Fission Yeast Fbh1. PLoS Genet. 10.

Unk, I., Hajdu, I., Blastyak, A., and Haracska, L. (2010). Role of yeast Rad5 and its human orthologs, HLTF and SHPRH in DNA damage tolerance. DNA Repair (Amst). 9, 257–267.

Uzunova, K., Göttsche, K., Miteva, M., Weisshaar, S.R., Glanemann, C., Schnellhardt, M., Niessen, M., Scheel, H., Hofmann, K., Johnson, E.S., et al. (2007). Ubiquitin-dependent proteolytic control of SUMO conjugates. J. Biol. Chem. 282, 34167–34175.

Vujanovic, M., Krietsch, J., Raso, M.C., Terraneo, N., Zellweger, R., Schmid, J.A., Taglialatela, A., Huang, J.W., Holland, C.L., Zwicky, K., et al. (2017). Replication Fork Slowing and Reversal upon DNA Damage Require PCNA Polyubiquitination and ZRANB3 DNA Translocase Activity. Mol. Cell 67, 882-890.e5.

Wright, W.D., and Heyer, W.-D. (2014). Rad54 Functions as a Heteroduplex DNA Pump Modulated by Its DNA Substrates and Rad51 during D Loop Formation. Mol. Cell.

Yuan, J., and Chen, J. (2011). The role of the human SWI5-MEI5 complex in homologous recombination repair. J. Biol. Chem. 286, 9888–9893.

Zellweger, R., Dalcher, D., Mutreja, K., Berti, M., Schmid, J.A., Herrador, R., Vindigni, A., and Lopes, M. (2015). Rad51-mediated replication fork reversal is a global response to genotoxic treatments in human cells. J. Cell Biol. 208, 563–579.

