## Supplementary materials for "Rrp1 translocase and ubiquitin ligase activities restrict the genome destabilising effects of Rad51 in fission yeast"

**Table S1** Strains used in this study

| Strain | Genotype | Reference |
|---|---|---|
| YA254 (WT) | ura4-D18, leu1-32, his3-D1, arg3-D1, h90 | (Akamatsu et al., 2003) |
| *rrp1Δ* | rrp1D::natMX, his3-D1, leu1-32, ura4-D18, arg3-D1, h90 | lab stock |
| *rrp2Δ* | rrp2D::natMX6, his3-D1, leu1-32, ura4-D18, arg3-D1, h90 | lab stock |
| SP282 *nmtP3-GFP-rad51* | kan-nmtP3-GFP-Rad51, leu1-32, ura4-D18, h- | a |
| *nmtP3-GFP-rad51* | kan-nmtP3-GFP-Rad51, his3-D1, leu1-32, ura4-D18, arg3-D1, h- | b |
| *nmtP3-GFP-rad51 rrp1Δ* | kan-nmtP3-GFP-Rad51, rrp1D::kanMX6, his3-D1, leu1-32, ura4-D18, arg3-D1, h- | b |
| *nmtP3-GFP-rad51 rrp2Δ* | kan-nmtP3-GFP-Rad51, rrp2D::kanMX6, his3-D1, leu1-32, ura4-D18, arg3-D1, h- | b |
| *nmtP3-GFP-rad51 p42-rrp2-HA* | kan-nmtP3-GFP-Rad51, ars1(MluI)::pREP42-HA-Rrp2 ura4+, his3-D1, leu1-32, ura4-D18, arg3-D1, h- | b |
| *254 p81-rrp1-FLAG* | ars1(MluI)::pREP81-Rrp1-FLAG, ura4+, his3-D1, leu1-32, ura4-D18, arg3-D1, h90 | b |
| *nmtP3-GFP-rad51 sfr1Δ* | kan-nmtP3-GFP-Rad51, sfr1D::arg3, arg3-D4, ura4-D18, leu1-32, h- | b |
| *nmtP3-GFP-rad51 swi5Δ* | kan-nmtP3-GFP-Rad51, swi5D::his3+, ura4-D18, leu1-32, his3-D1, arg3-D1, h- | b |
| *S. cerevisiae* AH109 | MATa, trp1-901, leu2-3, 112, ura3-52, his3-200, gal4D, gal80D, LYS2::GAL1UAS-GAL1TATA HIS3, GAL2UASGAL2TATA-ADE2, URA3::MEL1UAS-MEL1TATA-lacZ | CLONTECH |
| *E. coli Rosetta(DE3)* | *F- ompT hsdSB(rB- mB-) gal dcm (DE3) pRARE (CamR)* | Novagen |

a - from Hiroshi Iwasaki laboratory
b - this work

**Table S2** Plasmids used in this study

| Plasmid | Reference |
|---|---|
| pREP42-HA | (Craven et al., 1998) |
| pREP41-HA-Rrp1 | a |
| pREP41-HA-Rrp2 | a |
| pREP41-mCherry | a |
| pREP41-mCherry -Rrp1 | a |
| pREP81-FLAG | a |
| pREP41-Rrp1-FLAG | a |
| pREP81-Rrp1-DAEA-FLAG | a |
| pREP81-Rrp1-CS-FLAG | a |
| pREP81-Rrp2-FLAG | a |
| pREP42-EGFP | (Craven et al., 1998) |
| pREP42-EGFP-Rrp1 | a |
| pGADT7 | a |
| pGADT7-Rrp1 | a |
| pGADT7-Rrp2 | a |
| pGADT7-Rrp1 (1-226) | b |
| pGADT7-Rrp1 (227-482) | b |
| pGADT7-Rrp1 (483-694) | b |
| pGADT7-Rrp1 (695-897) | b |
| pGADT7-Rrp2 | a |
| pGADT7-Rad51N (1-117) | c |
| pGADT7-Rad51C (114-365) | c |
| pGBKT7 | a |
| pGBKT7-Rrp1 | a |
| pGBKT7-Rrp2 | a |
| pGBKT7-Rad51N (1-117) | c |
| pGBKT7-Rad51C (114-365) | c |
| pYK788 pREP1- $His_6$-Ubi1 | c |
| pREP41-RAD51 | c |

a - laboratory stock
b - this study
c - from Hiroshi Iwasaki laboratory

**Table S3** Primers used in this study:

| Cloned gene | Primer name | Primer sequence |
|---|---|---|
| FLAG | REP3flag_a_r | CTTTATCATCGTCGTCCTTGTAGTCGGATCCTCTAGAGTCGACATATGATTTAAC |
| | REP3flag_b_f | TACAAGGACGACGATGATAAAGACTACAAGGACGACGATGATAAAGACTA |
| | REP3flag_c_r | GGGTCATTTATCATCGTCGTCCTTGTAGTCTTTATCATCGTCGTCCTTGTAGT |
| | REP3flag_d_f | CAAGGACGACGATGATAAATGACCCGGGTAAAAGGAATGTCTCCCTTGCCAGTAC |
| Rrp1-FLAG | Rrp1_fwd | CAACTAATTATTCGAAACGGAATTCGAAACGATGGATTCATTGTCTGCATATC |
| | Rrp1_rev | TTTAAATGGCCGGCCGGTACCTCATGAATTAAGCCCAAATAG |
| Rrp1-DAEA-FLAG (D397A; E398A) | Rrp1_fwd | CAACTAATTATTCGAAACGGAATTCGAAACGATGGATTCATTGTCTGCATATC |
| | Rrp1D397A_rev | TATGTGCGGCCGCTAGAACAATGCGATAC |
| | Rrp1E398A_fwd | TGTTCTAGCGGCCGCACATACCATTCGT |
| | Rrp1E398A_rev | TTTAAATGGCCGGCCGGTACCTCATGAATTAAGCCCAAATAGATATAG |
| Rrp1-CS-FLAG (C609S) | R1c609sN_fwd | TTTCTGACTTATAGTCGCTTTGTTAAATCATATGGATTCATTGTCTGCATATC |
| | R1c609sN_rev | CAAACAAGGATCTAGACTAACACTACAGTTGAAATCC |
| | R1c609sC_fwd | AACTGTAGTGTTAGTCTAGATCCTTGTTTGGCTC |
| | R1c609sC_rev | TAGTCTTTATCATCGTCGTCCTTGTAGTCGGATCCTGAATTAAGCCCAAATAGATATAG |
| Rrp2-FLAG | Rrp2_fwd | CAACTAATTATTCGAAACGGAATTCGAAACGATGAGAAATAATACAGCTTTTGAAC |
| | Rrp2_rev | TTTAAATGGCCGGCCGGTACCTTATCGTGATGACATTCCAAATAAAAATG |
| Rrp1(1-226) | Rrp1_D1_fwd | ATAGTCGCTTTGTTAAATCATATGGATTCATTGTCTGCATATC |
| | Rrp1_D1_rev | CATCGTCGTCCTTGTAGTCGGATCCGGGAGTATTATGCTGAAG |
| Rrp1(227-482) | Rrp1_D2_fwd | ATAGTCGCTTTGTTAAATCATATGAGTCCGTTCGACACGATC |
| | Rrp1_D2_rev | CATCGTCGTCCTTGTAGTCGGATCCAGCCAAAAGGATGCGAAG |
| Rrp1(483-694) | Rrp1_D3_fwd | ATAGTCGCTTTGTTAAATCATATGTCTACGGTTTTTCGTAGAAC |
| | Rrp1_D3_rev | CATCGTCGTCCTTGTAGTCGGATCCTTCTTGTTCTGAAAAAGATTG |
| Rrp1(695-897) | Rrp1_D4_fwd | ATAGTCGCTTTGTTAAATCATATGAGCATTAAATTAAGGTGGG |
| | Rrp1_D4_rev | CATCGTCGTCCTTGTAGTCGGATCCTGAATTAAGCCCAAATAGATATAG |
| GST-Rrp1-FLAG, GST-Rrp1-CS-FLAG | Rrp1_fwd | GGGGCCCCTGGGATCTCATATGGATTCATTGTCTGCATATCC |
| | Rrp1_rev | CTCGAGTCGACCCGGGGATCCGCAAGGGAGACATTCCTTTTACC |

### Suplementary Figures

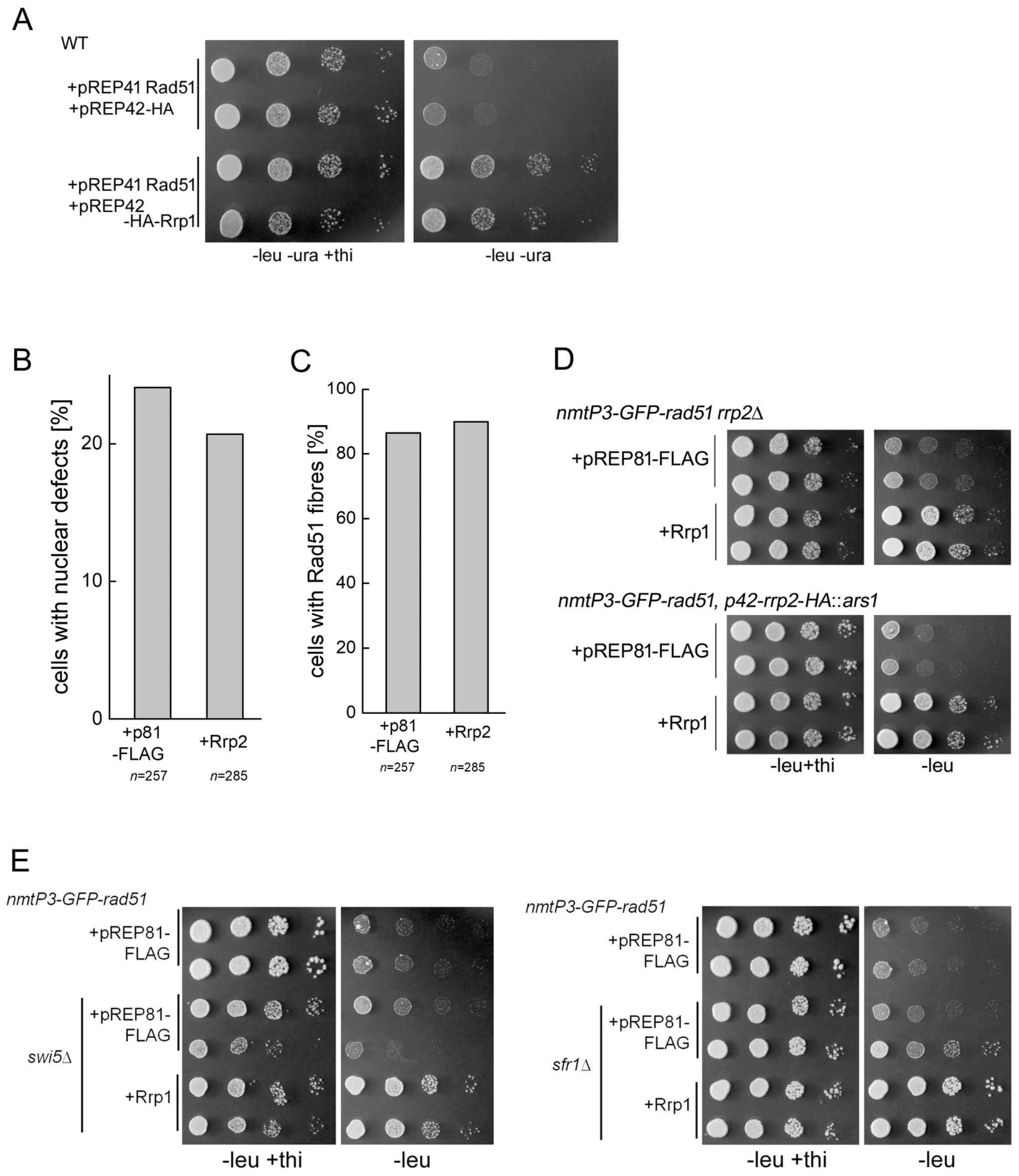


**Figure S1 Involvement of other factors in the Rad51-Rrp1 interaction**

(A) GFP tag is not responsible for the growth defect seen in GFP-Rad51 overproducing cells. Loss of viability is observed as growth inhibition on plates lacking thiamine (expression induction conditions). This is recused by overexpression of Rrp1. (B) Over-expression of *rrp2*$^+$ does not reduce the number of *nmtP3-GFP-rad51* cells undergoing aberrant mitosis. Cells with unequally segregated genetic material (cut and non-disjunction) were observed by DAPI staining of the nuclei of transformants grown for 48h in media lacking thiamine. n = total number of cells counted for 3 independent transformants for vector and *rrp2*$^+$. (C) Rad51 fibres are still present on chromatin in *nmtP3-GFP-rad51* cells overexpressing *rrp2*$^+$. n = total number of cells counted for 3 independent transformants for vector and *rrp2*$^+$ grown for 48 h in media lacking thiamine. (D) Deletion or overexpression of *rrp2*$^+$ does not affect suppression of the *nmtP3-GFP-rad51* growth defect by overproduction of Rrp1. (E) The rescue of the *nmtP3-GFP-rad51* growth defect by *rrp1*$^+$ overexpression is not dependent on the presence of *swi5*$^+$ or *sfr1*$^+$. Cells from cultures of two independent transformant were spotted for each strain in (A,D,E).

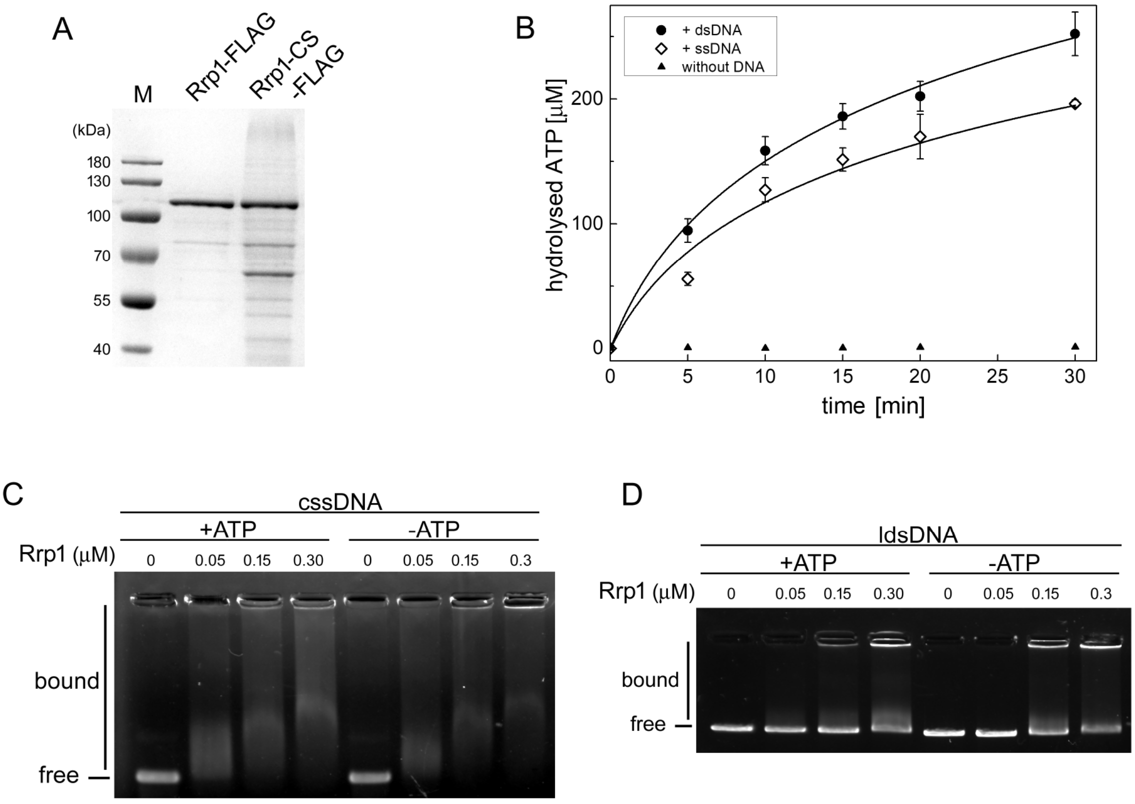


**Figure S2 Purified Rrp1 possesses DNA-dependent ATPase activity and binds both ssDNA and dsDNA.**

(A) Purified recombinant Rrp1 protein and a variant with mutated RING domain (Rrp1-CS) were analysed by SDS-PAGE followed by Coomassie Brilliant Blue staining. (B) Rrp1 has a robust DNA-dependent ATPase activity. (C) Electrophoretic mobility-shift assay (EMSA) demonstrating that Rrp1 binds to circular single-stranded DNA (cssDNA) in an ATP-independent manner. (D) EMSA demonstrating that Rrp1 binds to linear double-stranded DNA (ldsDNA) independently of ATP.

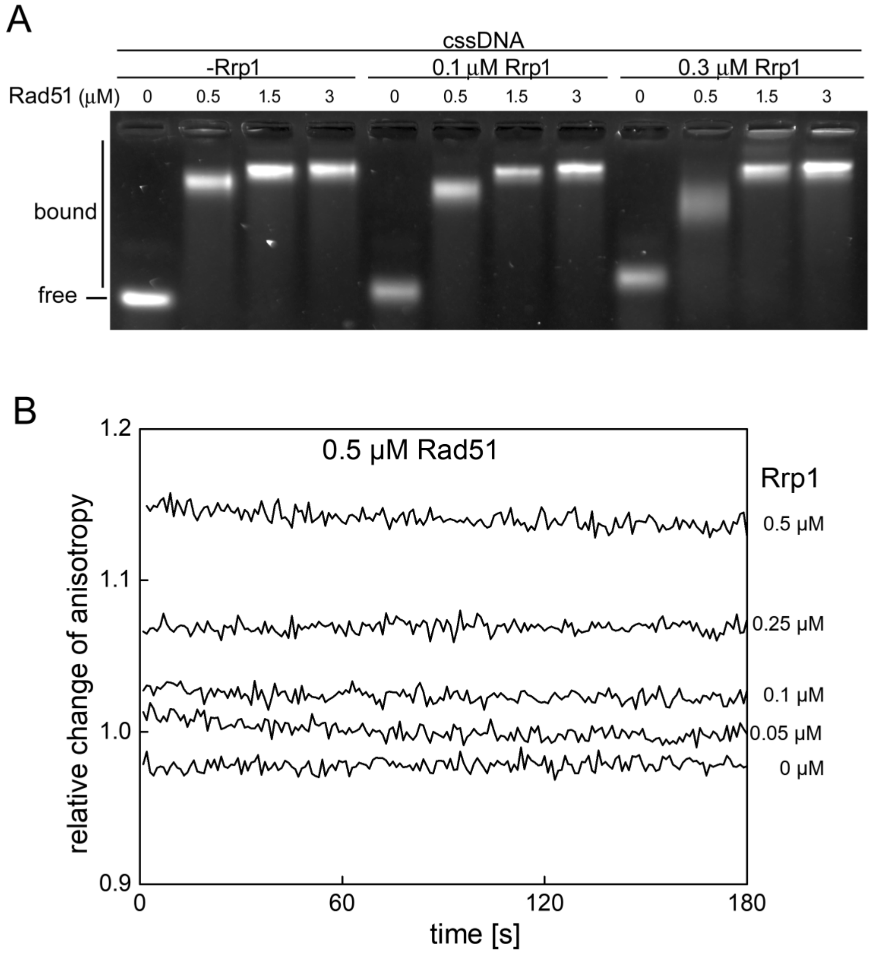


**Figure S3 Rrp1 does not disassemble Rad51-ssDNA complexes**

(A) Rrp1 has little effect on preformed Rad51-ssDNA filaments, as determined by EMSA. The indicated concentrations of Rad51 were incubated with cssDNA before the addition of Rrp1. (B) Rad51 filaments were formed on fluorescently labelled ssDNA, the indicated concentration of Rrp1 was added, and fluorescent anisotropy was measured in real-time. The addition of Rrp1 led to an increase, not a decrease, in relative anisotropy, indicating that Rrp1 does not disrupt Rad51-ssDNA complexes. Observed anisotropy increase is likely due to binding of Rrp1 to Rad51-ssDNA.

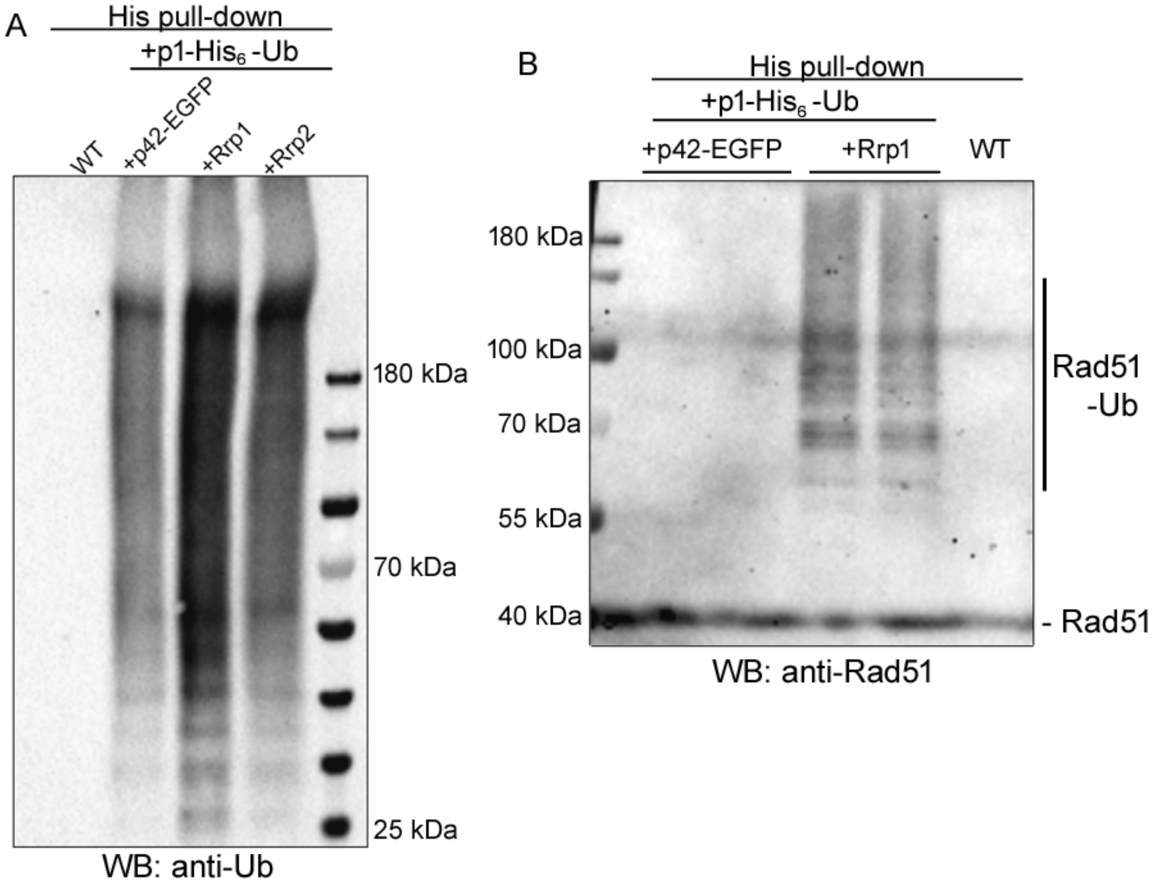


**Figure S4 Ubiquitylated proteins accumulate in cells overproducing Rrp1**

(A) Rrp1 overproduction leads to accumulation of ubiquitin modified proteins. The wild-type strain (WT) was co-transformed with a pREP1-based plasmid encoding hexahisitidine-tagged ubiquitin (+p1-$His_6$-Ub) together with a pREP42-EGFP-based plasmid either without an insert (+p42-EGFP), with Rrp1 (+Rrp1), or with Rrp2 (+Rrp2) . After 24 h growth in minimal media lacking thiamine (expression inducing conditions), ubiquitylated proteins were isolated by His pull-down and detected by western blot with an anti-Ubiquitin antibody. (B) Rrp1 overproduction results in the accumulation of ubiquitylated forms of Rad51. The experiment was performed as in (A) but Rad51 antiserum was used for western blot.
